# Two-timepoint assays of neural responses increase the sensitivity and specificity of single-cell whole-brain activity screens

**DOI:** 10.1101/2025.03.07.641687

**Authors:** Alejandro Ramirez, Lydia Rogerson, Evan J. Kyzar, Chloé Berland, Erica Rodriguez, Juan Guerrero, Luke Hammond, Anthony W. Ferrante, C. Daniel Salzman

**Author notes:** Department of Neurology, Ohio State University Medical Center, Columbus, OH, USA.

## Abstract

Current approaches for surveying whole brains for neurons activated during a particular state typically rely on immediate early gene (IEG) expression. However, IEG expression is variable across subjects and brain areas, demanding large sample sizes. Further, it cannot determine if the same or different neurons respond to two events. To overcome these issues, we present a whole-brain screening method utilizing transgenic mice to label neurons activated at two timepoints. An imaging and analysis pipeline surveys activity in ∼500 brain areas in different conditions. Compared to IEG methods, this approach reduces required sample sizes and enhances sensitivity and specificity. Finally, graph theoretical analyses are utilized to identify key circuit nodes - brain areas whose activity correlates with activity in other areas in a state-dependent manner. We validate this method by surveying whole-brain activity during hunger and satiety, and by investigating neural circuits activated by the GLP1 agonist semaglutide used to treat obesity.

## Introduction

In the last few decades, new microscopy methods have been developed that allow for high throughput imaging across a whole rodent brain at single-cell resolution (Ragan et al., 2012; Seiriki et al., 2017; Stelzer, 2015; Ueda et al., 2020). When these methods are applied in conjunction with staining for activity-dependent markers - such as expression of the immediate early gene (IEG) *c-Fos -* patterns of neural activation can be observed and quantified (Furth et al., 2018; Osten and Margrie, 2013; Parra-Damas and Saura, 2020; Seiriki et al., 2017; Tyson and Margrie, 2022). These approaches have been applied to study whole-brain neuronal activation during mating (Kim et al., 2015), thirst (Allen et al., 2019), the presentation of appetitive and aversive stimuli (Beyeler et al., 2018), and aspects of feeding behavior (Hrvatin et al., 2020; Nectow et al., 2017). Although the temporal specificity of IEGs is poor compared to calcium imaging or multi-unit recordings, it remains the only way to measure cellular activity across an entire mouse brain in relation to experimentally-controlled events.

Screening for activated neurons across the entire brain poses significant experimental and statistical challenges. In particular, despite the advances in imaging techniques, variability in neural activity among animals and the presence of noise can make it difficult to distinguish meaningful signals from background fluctuations, which in turn can dramatically limit the power of whole-brain activity screening. Adding to this, an assay for activity in relation to a single event, as is done in screening for IEG expression, cannot detect intermingled populations of neurons that respond to different stimuli. Indeed, using IEG staining alone allows one to assess in each mouse how the brain responds during only one event. In this scenario, when distinct populations of neurons in the same brain area are activated by opposing events, studies using IEG expression alone will not detect differences in the average activity across conditions (Logothetis, 2008; Stark et al., 2006). Consequently, the promise that these approaches can unbiasedly identify regions implicated in specific behaviors or brain states has been difficult to realize (Lara Aparicio et al., 2022; McReynolds et al., 2018; Phalip et al., 2024). In this paper, we develop an approach to whole-brain screening in mice that has enhanced statistical power compared to traditional IEG methods, and that also can detect areas with intermingled networks involved in different behaviors or internal states.

Traditional screening approaches that rely on IEG expression usually compare c-Fos counts of an experimental group (e.g., during a task state) to those of a control group (e.g., during a spontaneous state). The basic premise underlying c-Fos screening is that when neurons in a network or brain area are driven by a stimulus, their activity will increase, and this increased activity will be reflected in increased c-Fos expression. Therefore, comparing the number of neurons in a brain area that express c-Fos at baseline (i.e. in a control state) to an experimental state (e.g. a state evoked by a stimulus) will reveal regions specifically engaged by an experimental manipulation. However, the sensitivity and specificity of this approach is limited by the fact that not all neurons responsive to a stimulus will express c-Fos with each trial or manipulation (false negatives), and because spontaneously active neurons (neurons not involved in the stimulus or manipulation) also can express c-Fos (false positives). Therefore, c-Fos screening often requires many samples within both experimental and control groups to have sufficient power to identify neuronal populations of interest (Allen et al., 2019; Richman et al., 2023; Terstege and Epp, 2022). For whole brain screens, even more samples are required to detect activity driven by the event of interest due to the need for multiple comparisons corrections (Bennett et al., 2009). Sensitivity and specificity can be further compromised if a given region contains two distinct but intermingled neuronal populations active – one active in the control state and the other in an experimental state. In this circumstance, the control and experimental states both elicit c-Fos expression, obscuring the important qualitative differences in activity in each state.

In recent years, it has become possible to use IEG expression to indelibly label activated neurons, allowing for activity-dependent labeling in mice in at least two separate timepoints (Das et al., 2023; DeNardo and Luo, 2017; DeNardo et al., 2019; Gore et al., 2015; Sha et al., 2020; Shi et al., 2024). In a first timepoint, the promotor for an IEG drives the expression of an indelible label (e.g. a fluorescent protein) which can be visualized much later. Activity at the second timepoint is visualized by virtue of antibody staining for IEG expression itself. These tools have been used as a way to study engrams, or how neural representations change over time (Gore et al., 2015; Jung et al., 2023; Mayford and Reijmers, 2015; Roy et al., 2022), but not as a way of increasing the sensitivity and specificity of IEG screens or for detecting functional networks. We reasoned that the analysis of temporally separated events *within the same animal* could be useful for screening activated brain areas and allow for more robust whole-brain activity screens. We apply this experimental approach in different groups of mice tested in either the same or different conditions at each timepoint. We then employ statistical analyses to identify brain areas activated more (or less) than expected by chance by a particular event.

A strength of a two-timepoint approach is that it can be used to identify brain areas containing distinct populations of neurons, each activated by different events. For example, the amygdala has been shown to respond to both appetitive and aversive stimuli with distinct populations of neurons that are found within the same brain area (Gore et al., 2015). In an experiment where mice are studied using appetitive stimuli to drive one set of neurons, and aversive stimuli to drive another set of neurons, single timepoint screening using IEGs cannot distinguish whether the same or different neurons in any brain area are being activated. However, by using a two-timepoint approach, one can determine if each of the two events significantly activates neurons in a brain area, and whether these activated neurons are the same or different.

In this paper, we validate our whole-brain screening approach by comparing activity in two opposing states: hunger and satiety. We chose these states because substantial progress has been made in characterizing sensory, motor, and reward circuits in the brain that respond to caloric deficit or satiation (Andermann and Lowell, 2017; Morton et al., 2006; Morton et al., 2014). Using just five animals per experimental group and looking across ∼500 brain areas, we identified, in an unbiased manner, a majority of brain areas known to be modulated by fasting or refeeding. Furthermore, we identified brain areas containing spatially intermingled neurons differentially activated in the two states, such as the arcuate nucleus of the hypothalamus (ARH). Finally, we use our approach to identify brain areas activated by the hunger hormone ghrelin and the appetite-reducing medication semaglutide. After identifying these activated brain areas, we applied graph theoretical analysis to identify key nodes on the basis of the degree to which the number of activated neurons in one brain area correlates (or anti-correlates) with the number of activated neurons in other brain areas, in a state-dependent manner. Our unbiased whole-brain approach can therefore identify brain areas activated by multiple states or functions. It also identifies brain areas that could be important nodes for modulating neural circuits in a state-dependent manner to implement a particular behavioral function. Our approach offers significant promise for discovering distributed networks responsible for a variety of coordinated brain functions and behavioral states.

## Results

### Two-timepoint (TTP) Analysis Improves Sensitivity and Specificity of IEG Screens

To improve the sensitivity and specificity of whole-brain screening using c-Fos, we developed an assay that uses double-labeling to detect whether a neuron is activated by the same stimulus or condition at two distinct timepoints. We reasoned that this requirement can reduce the impact of non-specific, background activity of neurons, since double-labeling is less likely to occur simply due to chance. Moreover, this result can be compared to groups of animals who do not receive the stimulus, which provides a baseline assessment of the number of double-labeled cells (cells active at both time points) in the absence of the experimental manipulation. Assuming independent and uniformly distributed non-specific c-Fos expression or noise in each brain area, the probability of observing double-labeled cells can be analytically determined. This allows a *p*-value to be calculated that describes the probability that the experimentally observed number of double-labeled cells was greater than or less than chance (**Figure 1C, see Methods**). This scenario is equivalent to sampling “without replacement” and is described by a hypergeometric distribution, where ‘N’ is the total cell count of the brain area, ‘K’ is the number of cells labeled at timepoint 1 (TP1),‘n’ is the number of cells labeled at timepoint 2 (TP2), and ‘k’ is the number cells labeled at both timepoints. The hypergeometric distribution provides a null distribution for hypothesis testing concerning the number of neurons observed as labeled at both timepoints.

Using these analytic methods, a brain area is identified as being activated by a stimulus if the same stimulus is delivered twice and the observed number of cells activated at both timepoints is greater than expected by chance. Conversely, if a stimulus is not driving the neural activity, then the observed number of double-labeled cells should be indistinguishable from chance activation (i.e. indistinguishable from the null distribution). Note that this method allows *p*-values – describing whether the number of double-labeled cells is different from chance – to be calculated within individual animals. We then combine *p*-values across animals using a harmonic-mean approach (Wilson, 2019) that is robust to outliers and makes no assumption of independence. The resulting statistic is what we refer to as two-timepoint statistical inference (TTP_i_) (**Figure 1C**). A key requirement for calculating TTP_i_ is counting not only the cells labeled at each timepoint, but also the total number of cells in the area. The high density of cells within the mouse brain makes accurate quantification of all neurons challenging when using standard clearing and light-sheet microscopy techniques (Molbay et al., 2021), though recent advances have enabled more efficient imaging at a sufficient resolution and would allow the extension of our approach to those datasets.

To demonstrate the increased statistical power of TTP_i_, we simulated an experiment to detect differences in brain activity between a control condition and a stimulus-driven condition. We modeled a brain area with 5000 neurons each obeying Poisson statistics with a spontaneous average firing rate of λ. Within this network, a subnetwork of 250 neurons (5% of the total neurons) displayed a stimulus-driven firing at a rate of λ_s_. Across a number of different spontaneous (λ) and stimulus-evoked firing rates (λ_s_), we calculated the sample size needed for an adequately powered whole-brain study that relied on single-timepoint c-Fos counts (1TP, **Figure 1A**), two-timepoint raw counts of double-labeled cells (2TP, **Figure 1B**), or two-timepoint statistical inference, TTP_i_ (**Figure 1C**). When the background spontaneous activity level was set at λ=0.1 and stimulus-driven activity at λ_s_= 0.4, within the range of values commonly observed in electrophysiological data (de Kock and Sakmann, 2009; Ramirez et al., 2014), a traditional c-Fos*-*screen required 10 mice to detect differences between the task state and spontaneous state. By comparison, TTP_i_ achieved sufficient statistical power with only three animals (**Figure 1D**). TTP_i_ thereby significantly reduces the sample size requirements for informative whole-brain screening analysis.

**Figure 1.**
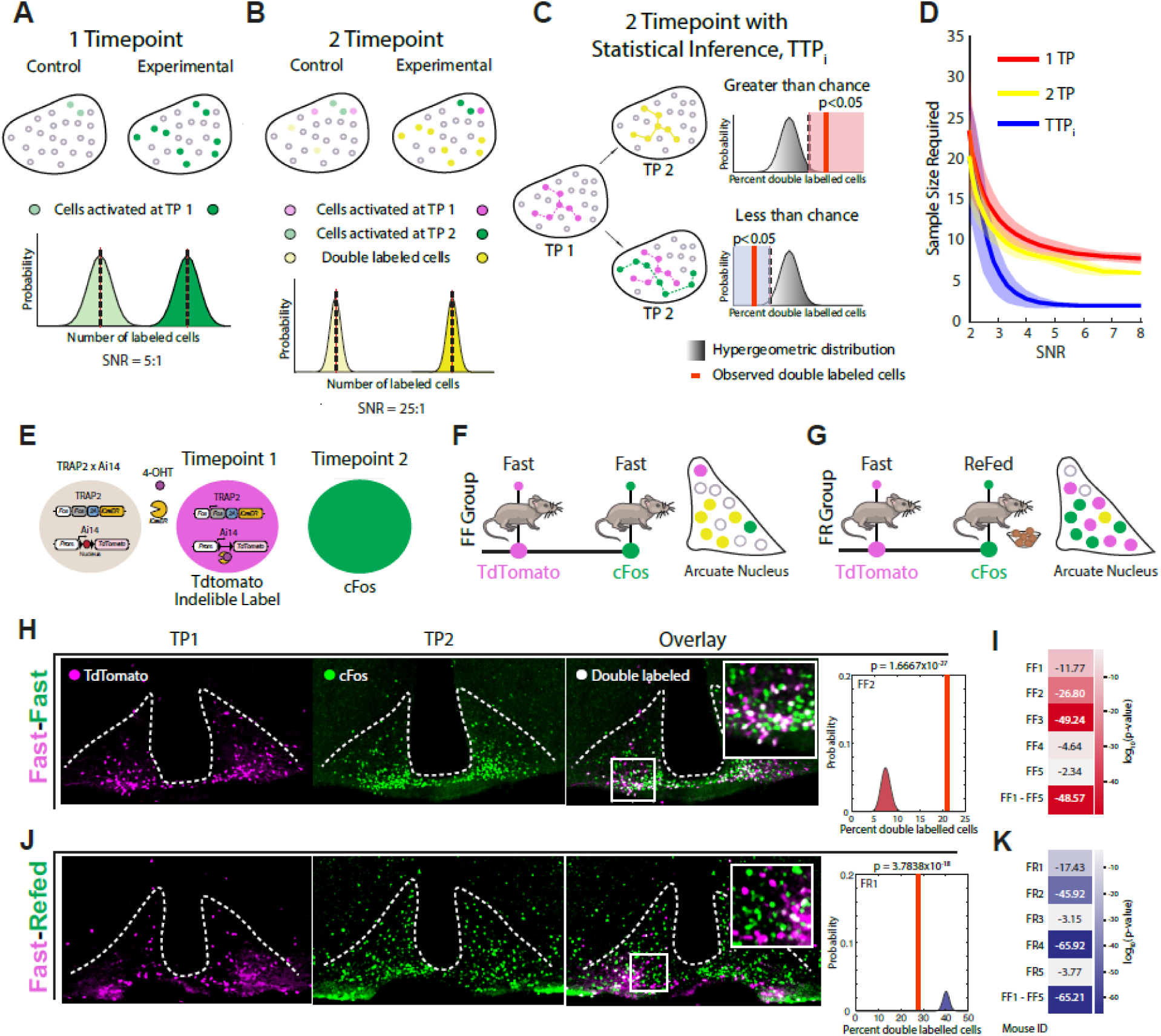
A Two-timepoint Labeling Improves Sensitivity of IEG Activity Screening. **A)** Schematic of traditional IEG screening approach relying on a single timepoint (1TP) such as c-Fos. The average c-Fos counts in a control group (spontaneous activity; 0.1 a.u. in this example) are compared to the c-Fos counts of an experimental group (task/stimulus activity; 0.5 a.u. in this example), and statistics are used to determine if samples are drawn from the same or different distributions. The signal-to-noise ratio in this hypothetical example is 5:1. **B)** A two-timepoint assay (2TP) allows comparison of double-labled cells across 2 independent timepoints in groups of mice, improving the signal-to-noise ratio (noise: 0.1 x 0.1 = 0.01; signal: 0.5 x 0.5 = 0.25; signal-to-noise = 25:1). **C)** TTP statistical inference (TTPi) approach for within-animal statistical inference, detecting whether the observed number of double-labeled cells (or lack thereof) is statistically significant within individual animals. A *p*-value falling on the right of probability distribution (shaded in red) indicates more double-labeled cells than expected, and a *p*-value on the left of the distribution (shaded in blue) indicates fewer double-labeled cells than expected. **D)** Power analysis showing sample size requirements for a whole-brain analysis using 1-timepoint (1TP; e.g., c-Fos) vs. 2TP vs. TTPi at different signal-to-noise ratios. Solid lines indicate mean and shaded areas represent standard error. **E)** An *in vivo* two-timepoint system was achieved by crossing TRAP2 mice with Ai14 reporter animals. This system labels cells active at timepoint 1 by virtue of tdTomato (tdT) expression. Active cells at timepoint 2 can be identified by staining for c-Fos, thus achieving two timepoint activity labeling. **F)** Fast-Fast (FF) TTP behavioral assay, labeling neurons which are active during both timepoints of fasting. **G)** Schematic of Fast-Refed (FR) TTP behavioral assay, which should detect less double-labeled cells than expected in brain regions such as the arcuate nucleus of the hypothalamus (ARH) in neurons activated by fasting and refeeding, respectively. **H)** A high-resolution image of the ARH in a representative FF mouse. Timepoint 1 (tdT: magenta), timepoint 2 (c-Fos: green), and the overlay with inset showing clearly double-labled cells. TTPi plot shown on the right for an example mouse (FF2 in Figure 1I) with significant *p*-value for overrepresentation compared to expected double-labled cells. (p=1.7x10^-27^). **I)** *P*-values for 5 individual FF animals and combined *p*-value (FF1-FF5) represented as a heatmap in red, all showing highly significant *p*-values. **J)** Same as D except for Fast-Refed (FR) brain, with inset showing low number of double-labeled cells activated by fasting and refeeding. TTPi plot shown on the right for an example mouse (FR1 in Figure 1K) with significantly fewer double-labeled cells than expected by chance; (p=3.8x10^-18^). **K)** *P*-values for 5 individual FR animals and combined *p*-value (FR1-FR5) represented as a heatmap in blue, all showing significant *p*-values for fewer double-labeled cells than expected.

### Two-timepoint Assay in Mice

We next sought to demonstrate that robust two-timepoint statistical inference could be successfully implemented in an *in-vivo* experiment across the whole brain. To perform biological two-timepoint assays in mice, we crossed transgenic TRAP2 animals (Allen et al., 2019; DeNardo et al., 2019) with Ai14 reporter animals (Madisen et al., 2010), yielding double transgenic TRAP2:Ai14 animals (**Figure 1E**). This inducible system, relying on expression of *Cre-ER* driven by the c-*Fos* promoter, allows for activity-dependent expression of the fluorescent reporter tdTomato (tdT) which indelibly labels active neurons at timepoint 1. Since TRAP2 animals retain endogenous *c-Fos* activity, subsequently staining the brain for c-Fos protein at timepoint 2 achieves labeling at two independent timepoints.

To test the TRAP system as a means of utilizing our two-timepoint statistical approach, we designed an assay to capture cells that are active during two opposing physiological states – hunger (fasting) and satiety (refeeding). We focused our initial analysis on activity in the ARH since this brain area is known to have distinct populations of neurons active under each of the two conditions. For timepoint 1, mice were given 4-hydroxytamoxifen (4-OHT) at the beginning of their dark cycle following an 18-hour fast to indelibly label neurons activated during hunger. Mice were then returned to the homecage for one month. After one month, mice were fasted for 22.5 hours. Half the animals continued to be fasted for 1.5 hours (i.e., fasted at both time points = Fast-Fast or FF) while the remaining were refed for 1.5 hours (i.e., fasted at the first time point and refed at the second = Fast-Refed or FR). Brains were collected and processed to detect tdT fluorescence and c-Fos protein expression. We predicted that FF animals would show more double-labeled cells than expected by chance in the ARH due to activity of putative AgRP “hunger” neurons at both timepoints, whereas FR animals would demonstrate fewer double-labeled cells than expected by chance due to the existence of reciprocally-activated AgRP and POMC neurons activated by fasting and refeeding, respectively (Mandelblat-Cerf et al., 2015) (**Figure 1F,G**).

Cells activated at both timepoints were readily identifiable in FF mice - TTP_i_ analysis of activity in the ARH was highly significant (*p*=1.7x10^-27^) (**Figure 1H**). Meanwhile, in the FR condition, our TTP_i_ analysis indicated there was a significant lack of double-labeling of cells activated at the two timepoints, consistent with prior evidence indicating that the ARH contains distinct populations of cells active during hunger and satiety (*p*=3.8x10^-18^, **Figure 1J**). These results were consistent across five experimental subjects in each group (**Figure 1I,K**). Control experiments demonstrated that reversing the order of fasting and refeeding at the two timepoints yields the same results (**Figure S1A**). In addition, measuring activation of neurons in the ARH at two timepoints in *ad-libitum* fed mice (mice that were neither fasted nor refed after fasting) in the homecage did not result in double-labeling of more neurons than would be expected by chance (**Figure S1B**). Thus, the TTP_i_ method can detect significant activation of neurons in a region of the brain known to be responsive to feeding and fasting, and the low *p*-values suggest the potential use for TTP_i_ for screening whole-brain activity.

### Whole-Brain Imaging and TTP Analysis Pipeline

To apply the TTP approach across the whole brain, we developed a pipeline with a number of features often difficult to achieve using a single methodology: 1) the ability to register the brain to a common coordinate framework, 2) the ability to restore low signal-to-noise data for improved segmentation accuracy, 3) the ability to detect labeled cells across the brain, 4) the ability to detect co-labeled cells activated at two timepoints, and 5) the ability to determine the total cell count in each surveyed brain area. Methods that rely on clearing the brain, such as iDISCO and ClearMap, are excellent for high-throughput imaging of whole brains and for counting single-channels (e.g. for c-Fos mapping); however, efficiently imaging at sufficient resolution to accurately quantify areas of dense neuronal populations presents significant challenges, which can be compounded by difficulties obtaining consistent antibody penetration (Molbay et al., 2021). As TTP_i_ requires total neuronal cell counts for each brain area examined, we utilized serially sectioned mouse brain tissue and refined an open-source Fiji software plugin for whole-brain registration and analysis called BrainJ (Botta et al., 2020). TRAP mice experienced two events of interest (e.g. fasted, refed) separated by one month. At the time of the first event, mice were treated with tamoxifen. After the last event, the entire brain from each mouse was fixed, sectioned on a cryostat, stained for c-Fos protein, and imaged using spinning disk confocal microscopy to detect both c-Fos and tdT dependent fluorescence and neurons. Each brain was then analyzed using our refined Python and Fiji based BrainJ software that registers sections to the Allen Brain Atlas Common Coordinate Framework (**Figure 2A,B**). To improve the accuracy of cell detection in lower resolution images with a reduced signal-to-noise ratio, we trained a Content Aware Image Restoration Network (CARE) to improve image quality and trained convolutional neural network-based U-Nets for automated cell segmentation to obtain total cell counts (Chen et al., 2021) (**Figure 2C**).

Our approach successfully provided quantification of four parameters – tdT count (K), c-Fos count (n), total cell count (N; which can be either NeuN or DAPI counts – see Methods), and the number of double-labeled cells by c-Fos and tdT (k) – that enabled us to apply TTP_i_ to each brain area registered with the Allen Brain Atlas. We then could visualize *p*-values across the brain (**Figure 2D,E; also see Figure S2**). Note that the whole-brain pipeline used a 4x 0.2 NA Plan Apochromat objective and leveraged deep learning image restoration and segmentation for cell detection. Despite the reduced resolution and signal obtained when compared to brain area-specific data acquired using a 10x 0.45 NA Plan Apochromat objective, our approach detected significant double-labeling of cells in the ARH of the same brains that were fasted at both time points (FF) (**Figure 2F**). This suggests that our protocols for registration, restoration, segmentation, quantification, and TTPi analysis could be applied across the whole brain and yield interpretable results. However, the whole-brain approach did not detect a significant lack of double-labeled cells in the ARH of the FR brain (**Figure 2G**), suggesting a limitation in the ability of our whole-brain approach to detect intermingled, functionally-opposed neuronal populations – likely due in part to decreased resolution in our whole-brain TTP_i_ pipeline.

**Figure 2.**
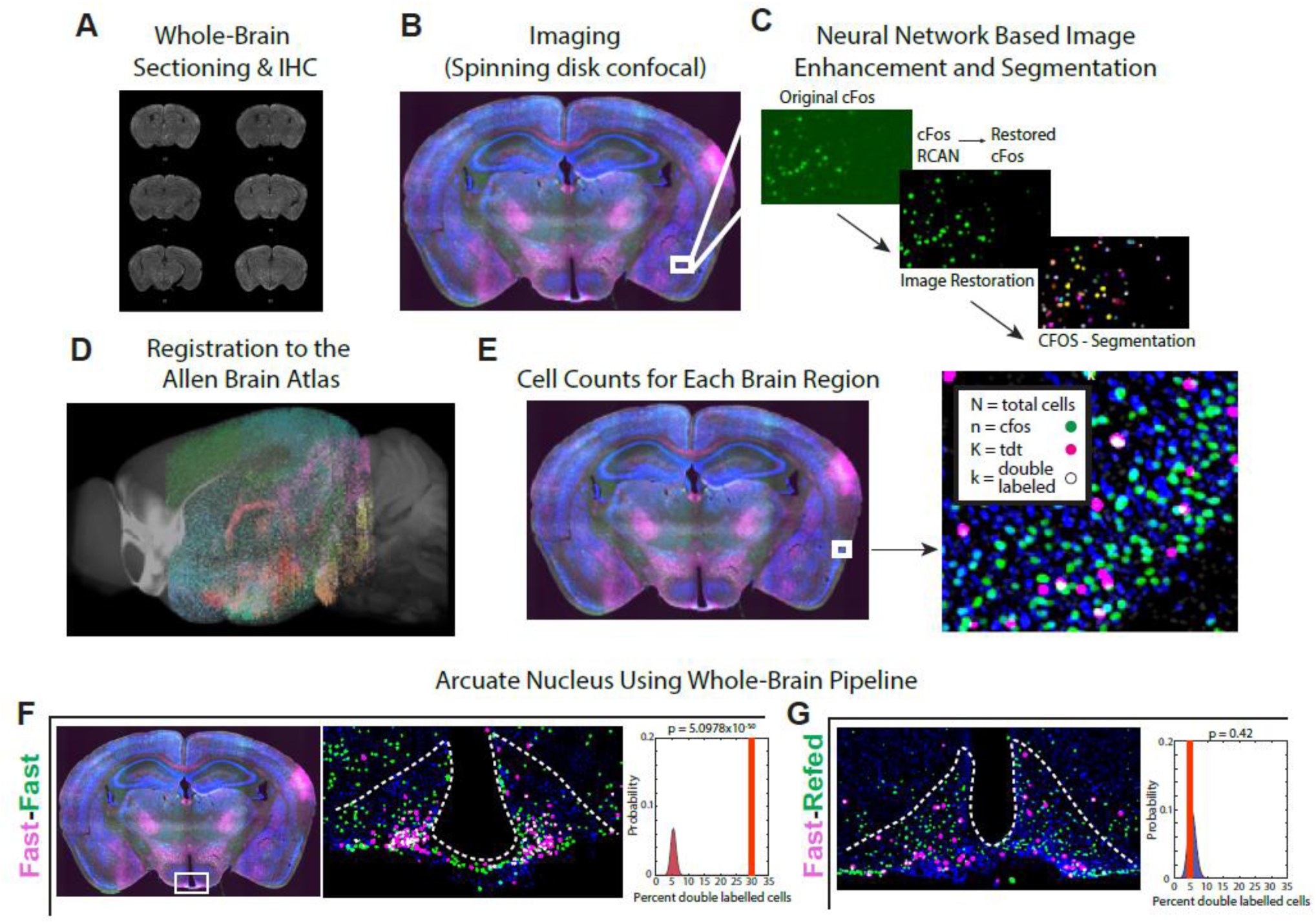
**Whole-brain pipeline for two-timepoint statistical inference (TTPi)**. **A)** Whole brains were sectioned at 50 microns, stained for c-Fos, NeuN, and DAPI, and then **B)** imaged using a spinning-disk confocal at 4x resolution**. C)** Images are pre-processed by training a Content Aware Image REstoration Neural Network (CARE) to enhance resolution of 4x images up to 10x resolution, allowing more accurate automated segmentation of cell counts. Note that while only the c-Fos channel is shown, this is done for all 4 channels (see Figure S2). **D)** Whole brains were then registered to the Allen Brain Atlas Common Coordinate Framework (CCF) using open-source software, BrainJ (Botta et al., 2020). The colored area of brain represents extent of antero-posterior registration for this example brain. **E)** A 50um^2^ section of tissue with cell-detection for total cell count (N; for which either NeuN or DAPI can be used – see Methods), c-Fos (n), tdt (K), and double-labled cells (k). These parameters are determined for every brain area corresponding to regions of the Allen Brain Atlas CCF, allowing *p*-values to be calculated for each region. **F)** Coronal section of Fast-Fast (FF) brain and segmentations for arcuate nucleus of the hypothalamus (ARH) (middle panel) along with TTP statistical inference (TTPi right), validating the ability for the whole-brain pipeline to determine if a brain region shows significantly more double-labled cells than expected. **G)** Same as H except showing segmentation (left panel) and TTPi (right panel) for Fast-Refed (FR)brain, indicating the limitations of TTPi in detecting significant under-representation of expected numbers of double-labeled cells.

### Simulations to Assess the Strengths and Limitations of TTP_i_

We next sought to evaluate the sensitivity of TTP_i_ for determining if the number of double-labeled cells was less than or more than expected by chance in the parameter space typical of our whole-brain data. To accomplish this, we performed a simulation whereby we varied the total cell count (N) and the proportion of cells activated at both timepoints and then computed a *p*-value using TTP_i_ for each simulated brain area (see Methods). This simulation assumed that, on average, a similar proportion of neurons was activated by a stimulus at each of the two timepoints. When alpha values were set to 0.0001 to account for multiple comparisons corrections needed for whole-brain analysis (i.e. Bonferroni correction – *p*=0.05/500), TTP_i_ provided high sensitivity for detecting double-labeled neurons (neurons active at both timepoints) across all conditions with the exception of brain areas where the total cell count was less than 5000 neurons and the proportion of activated cells was less than 5% of the total (in other words small brain areas with few active cells; **Figure 3A**). When the simulation results were overlaid on data parameters obtained from actual mouse brains (see Methods), 21,902 out of 22,451 brain areas analyzed across 40 brains, (97.2%) were within the range of sensitivity for detecting significant overrepresentation of overlapping cell labeling. Conversely, TTP_i_ tends to fail to detect a significant lack of double-labeled cells (i.e., high false negative rate) when the fraction of activated cells is low – as is typically observed for real data. In this case, small proportions of cells exhibiting activity at both timepoints fall within the null distribution computed from the hypergeometric distribution (see Methods). Under parameter ranges observed in our whole-brain data, the simulations were sensitive for detecting less double-labeling than expected by chance in only 5,210 out of 22,541 brain areas (23%) (**Figure 3B**).

TTP_i_ detects brain areas where the number of neurons activated at two timepoints is more than expected by chance. However, if a particular subpopulation of neurons within a brain area exhibits a higher basal firing rate than the remaining neurons, then TTP_i_ could inaccurately report that double-labeled cells are more frequent than expected by chance, even though this higher firing rate may be unrelated to the experimental manipulation being investigated. For example, if a subpopulation of neurons has a higher spontaneous firing rate than the rest, these neurons are more likely to be active spontaneously at both timepoints, resulting in a false positive when applying TTP_i_. We verified that TTP_i_ has a high false positive rate in these circumstances by performing a simulation where a 0.25-1.5% of all cells in a given brain area had a baseline firing rate (from 0.2 to 1) that was distinct from stimulus-responsive cells (where the firing rate was held at an average of 0.4). In this case, TTP_i_ has a propensity for finding that the number of double-labeled cells is greater than expected by chance, even though this activity was not driven by the stimulus or condition being investigated (a false positive, **Figure 3C** and **Figure S3A-C**).

### Improving the Sensitivity and Specificity of Whole-Brain Screening with TTP Analysis with Subtraction (TTP-S)

One way to reduce the false positive rate when trying to identify condition-specific neuronal activation is to perform TTP_i_ on a control group of mice which does not experience the same experimental manipulation, and then to compare those results with the experimental group. This comparison can, in essence, subtract the contribution of subpopulations of neurons with inherently higher firing rates that can lead to false positives with TTP_i_. Comparisons between different groups of mice may also help increase sensitivity for detecting significantly fewer double-labeled cells than expected by chance.

To achieve comparisons between groups of mice, we use a bootstrapping approach to compare hypergeometric distributions from two animals tested in different conditions along with the experimentally observed double-labeling count from the two animals. We sample repeatedly from the individual hypergeometric distributions computed for each mouse and take the difference between the samples to produce a new null distribution. We then compute a new *p*-value by asking how many samples in the distribution are equal to or greater, or equal to or less than, the difference of the experimentally observed double-labeling count. We call this approach TTP with subtraction (TTP-S). TTP-S is performed by sampling from all pairwise relationships across animals from the two groups (experimental vs. control). Each experimental animal is then tested for significantly over-or underrepresented double-labeled cells in each brain area, and a combined *p*-value is obtained across animals in a group by using a harmonic mean approach.

To gain an intuition for how this approach reduces false positives, consider the case when one is surveying the brain for areas where neurons are activated in an experimental condition (condition A) at each of two timepoints (AA mice). The effect of a subpopulation of neurons with a higher firing rate can be subtracted out by considering data from control mice where, e.g., condition A is tested at timepoint 1, and condition B at timepoint 2 (AB mice, **Figure 3D**). The assumption is that the subpopulation of neurons with a higher firing rate will be observed in both condition A and B. As a result, TTP-S subtracts their contribution to chance observation of double-labeled cells, thereby decreasing the false positive rate (**see Methods**, **Figure 3D**). Similarly, if one performs TTP-S using AB mice compared to mice in the homecage at each of timepoints (Homecage-Homecage or HH mice), the false negative rate can also be decreased (**Figure 3E**), as the probability of lack of double-labeling in the experimental AB group is being compared to the number of double-labeled cells in a control condition.

To compare the sensitivity and specificity of TTP-S to other methods for identifying differentially activated brain areas between experimental groups, we performed another simulation using parameter ranges that matched those from our experimental data, including the distribution of total cell counts (**Figure S3D**) and the ratio of activated cells (K/N ratios) (**Figure S3E**). The simulation evaluated each method’s ability to detect differences in brain activity between two states – state A and state B. In the simulation, 500 brain areas varied by size (1000 – 30,000 neurons), with each area containing subnetworks of neurons activated by one state, both states, or neither, as well as some areas with a tonically active subnetwork that differed in baseline activity from the other neurons. Of these 500 areas, 298 were stimulus-responsive (i.e. true positives for A and/or B). 116 of these areas had subnetworks functionally intermingled (i.e. responsive to both state A and state B). Further, 118 brain areas contained a tonically active subnetwork that was not stimulus-responsive. The background and stimulus-evoked firing rates were randomly assigned to produce a range of different signal-to-noise ratios. For each brain area, we simulated stimulus-evoked activity during state A and/or state B. Therefore, brain areas in the simulation had a wide variety of total cell counts, network sizes, spontaneous firing rates, and stimulus evoked firing rates.

After running the simulation, we compared the cell count differences in each brain area with simulations of one-timepoint activity in A vs B (meant to simulate the traditional screening method using only c-Fos), two-timepoint simulations looking at the differences in the raw number of double-labeled cells between AA mice and BB mice, TTP_i_ as described above (without subtraction) that can examine the significance of the number of double-labeled cells in individual AA or AB mice, and TTP-S for AA mice compared to AB mice (**Figure 3D**). We then compared the statistical results from these four methods using consistent sample sizes across all simulations (n=10, 5 AA ‘brains’ and 5 AB ‘brains’; **Figure 3F**). Receiver Operating Characteristic (ROC) curves were generated to evaluate the ability of the different statistical approaches – one-timepoint vs. two-timepoint vs. TTP_i_ vs. TTP-S – to distinguish between true and false positives across the different brain areas. The area under the ROC curve (AUC) provides a measure of the ability to correctly classify brain areas as responding to state A or state B.

Results from this simulation reveal that TTP-S achieves significantly improved sensitivity and specificity over traditional one-timepoint screening, with an AUC of 0.99 for TTP-S, 0.91 for TTP_i_, 0.92 for two-timepoint (2TP), and 0.58 for one-timepoint (1TP) (**Figure 3F**; see **Figure S3F** for confusion matrices).

**Figure 3.**
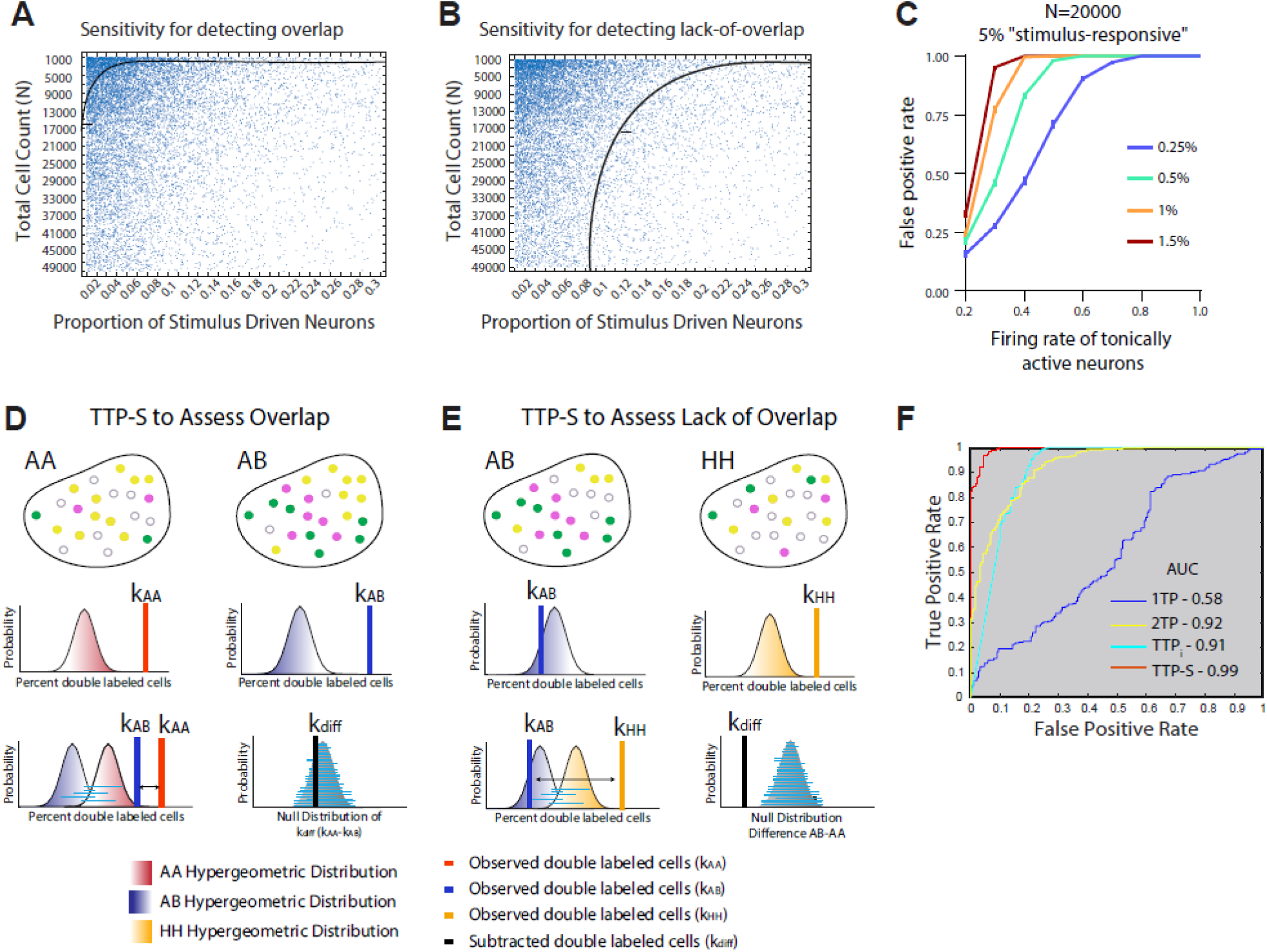
Limitations of whole-brain c-Fos screening are overcome with TTP with subtraction (TTP-S). We first evaluated the sensitivity for detecting overlap (more double-labeled cells than expected by chance) or lack of overlap (fewer double-labeled cells than expected by chance) in simulated brain areas using TTP with statistical inference (TTPi). The alpha value was set at *p*=0.0001 to account for multiple comparisons corrections needed for whole-brain analysis. We simulated a single brain area but varied the parameters of total cell count (N) and the proportion of active cells at two time points – K/N at timepoint 1 and n/N at timepoint 2. **A)** TTPi demonstrates high sensitivity for detecting significantly overrepresented numbers of double-labeled cells compared to chance. The black line indicates the cut-off point at *p*=0.0001 for detecting significantly higher numbers of double-labeled cells (all points to the right of the line). A scatter plot from our whole-brain analysis is shown on each plot, where each dot represents a single brain area. Out of 22,451 brain areas analyzed across 40 brains, 21,902 (97.2%) were within the range of sensitivity. **B)** TTPi demonstrates lower sensitivity for detecting fewer double-labeled cells than would be expected by chance (black line indicates cut-off point at *p*=0.0001, where all points to the right of the line show significantly less double-labeling), with only 5,210 out of 22,541 brain areas taken from our whole-brain analysis (23%) within the range of sensitivity. **C)** Simulated brain region with total cell count (N) of 20,000, of which 5% are stimulus-responsive at both timepoints at an average firing rate of 0.4. The firing rate of 0.25 – 1.5% of separate cells was varied from 0.2 to 1. Increasing the proportion of neurons in the tonically active network as well its firing rate increases the false positive rate. **D)** Schematic illustrating TTP with subtraction (TTP-S). By bootstrapping the difference between two null distributions, a new null distribution is created. The difference in double-labeled (overlapping) cells between AA and AB mice is compared to the new null distribution. The subtraction decreases the false positive rate observed if only using TTPi. **E)** Same as D), except showing how the TTP-S method also corrects for false negatives (fewer numbers of double-labeled cells than expected) seen if one only uses TTPi to detect lack of overlap in AB brains, with animals exposed to only the homecage environment at both timepoints (homecage-homecage or HH) used as a control group. **F)** Receiver operating characteristic (ROC) curves for correctly classifying true and false positives when using the 4 different screening approaches: one timepoint (1TP) vs. two timepoint (2TP) vs. TTPi vs. TTP-S in 500 simulated brain areas with varying total cell counts and stimulus-responsive populations. ROC curves measure the ability of the different approaches to correctly classify brain areas.

### Whole-Brain TTP Analysis with Subtraction Detects Brain Areas Activated by Fasting and Refeeding

More than two dozen brain areas in the mouse have been shown to respond to signals related to caloric deficit, incretin hormones and food intake (Andermann and Lowell, 2017; Morton et al., 2006; Morton et al., 2014). These prior studies provide experimental context to validate the ability of our whole-brain pipeline and TTP-S analysis to detect brain areas activated in response to experimental manipulations. As an initial test of the TTP-S pipeline, we therefore surveyed the whole brain for activated cells when mice were in two states: fasted (hungry) and refed (sated). After using our whole-brain pipeline to process and analyze brain images, we compared TTP-S to the other methods described. We created five groups of mice (n=5 each; n=4 in HH group), one for each combination of testing in the fasted and refed state at timepoints 1 and 2 (FF, FR, RR, F, and HH groups). For timepoint 1, FF and FR mice were ‘TRAPed’ with 4-OHT at the beginning of their dark cycle following an 18hr fast in order to label neurons that are active during fasting. They were returned to their homecage, and one month later, they either were fasted for 24 hours and then refed for 1.5 hours for Fast-Refed (FR) animals, or not refed for Fast-Fast animals (FF mice). In an analogous fashion, we created a Refed-Refed (RR) group, where mice were refed at both timepoints. We created an additional group in which both timepoints occurred in the control, homecage condition in the absence of any stimulus (HH mice), and finally a group where we only measured activity at one timepoint in the fasted state using c-Fos (F group) (**Figure 4A**). All mice in all groups were sacrificed after timepoint 2 (or after fasting in the F group), and whole brains were extracted and stained for c-Fos. Our study design allowed for a direct comparison of the different approaches and analyses for identifying brain areas activated in the fasted and refed states across the majority of the mouse brain.

We analyzed approximately 600 brain areas per animal (min = 570; max = 656), counting an average of 20 million DAPI+ cells (2.0014x10^7^ +/-1.1657 x10^6^); 13 million NeuN+ cells (1.34355 x10^7^ +/-1.2966 x10^6^); 1 million c-Fos+ cells (1.0765 x10^6^ +/-4.3767 x10^5^) and hundreds of thousands of tdT+ cells (2.3968 x10^5^ +/-9.9149 x10^4^) per mouse (**Figure S4A-C**). Note that these cell counts reveal that variability of TRAP expression (as indicated by tdT-positivity) was high, similar to prior studies (DeNardo et al., 2019), as was variability in c-Fos protein expression. Unfortunately, hindbrain areas were not consistently registered, and therefore were not included in our analyses.

We next compared methods for detecting differences between experimental groups. Classic one-timepoint brain assessment of activity compared anti-c-Fos staining in homecage controls (HH mice) to staining in FF and F or FR and RR brains for fasted and refed states, respectively (1TP). Two-timepoint assessment involved comparing the raw counts of double-labeled cells in FF and RR mice to HH controls (2TP). TTP_i_ was used in FF and RR brains. Finally, TTP-S used FF-FR brains and RR-FR brains, with the FR group being used for subtraction (**Figure 4A**).

Using a 1TP approach, we failed to detect significant differences in any brain area for either fasting or refed conditions compared to HH controls after correcting for multiple comparisons (**Figure 4B,C**). The 2TP method identified only one significant brain area for FF mice and six for RR mice compared to HH controls after correcting for multiple comparisons. Results were similar when comparing fasted and refed brains to each other rather than using HH mice as the comparison group (**Figure S4D,E**). By contrast, TTP-S identified 54 brain areas activated in the fasted condition and 108 brain areas in the refed condition that passed a more stringent significance level (see Methods). Note that TTP_i_ identified many more brain areas as being statistically different (**Figure 4B,C**), which could be due to the fact that this approach can have a higher false positive rate, as shown in our simulations (**Figure 3C**).

We next rank-ordered brain areas by the significance of *p*-values from TTP-S. In fasted brains, many well-known feeding-related brain areas appeared at the top of the list, including the ARH, the medial preoptic nucleus (MPO), and the dorsomedial hypothalamus (DMH) (Andermann and Lowell, 2017; Morton et al., 2014; Myers and Olson, 2012; Sternson and Eiselt, 2017) (**Supplemental File S1**). In refed brains, a number of brain areas previously implicated in the refed state were identified, including the paraventricular hypothalamic nucleus (PVH), tuberomammillary nucleus (TU), and parasubthalamic nucleus (PSTN). Notably, for both RR and FF brains, activity in the top 30 statistically significant brain areas tended to be activated in a correlated manner. To demonstrate this, we visualized the covariance of double-labeling across all brain areas using t-distributed stochastic neighbor embedding (t-SNE) analyses combined with a k-means cluster analysis. We then labeled the 30 brain areas with the lowest *p*-values according to TTP-S in the refed and fasted conditions (**Figure 4D,E**). Each of the 30 brain areas clustered together in both the fasted and refed brains, unlike the areas identified in FR TTP-S analysis or 30 randomly-chosen brain areas **(Figure S4F,G)**.

**Figure 4.**
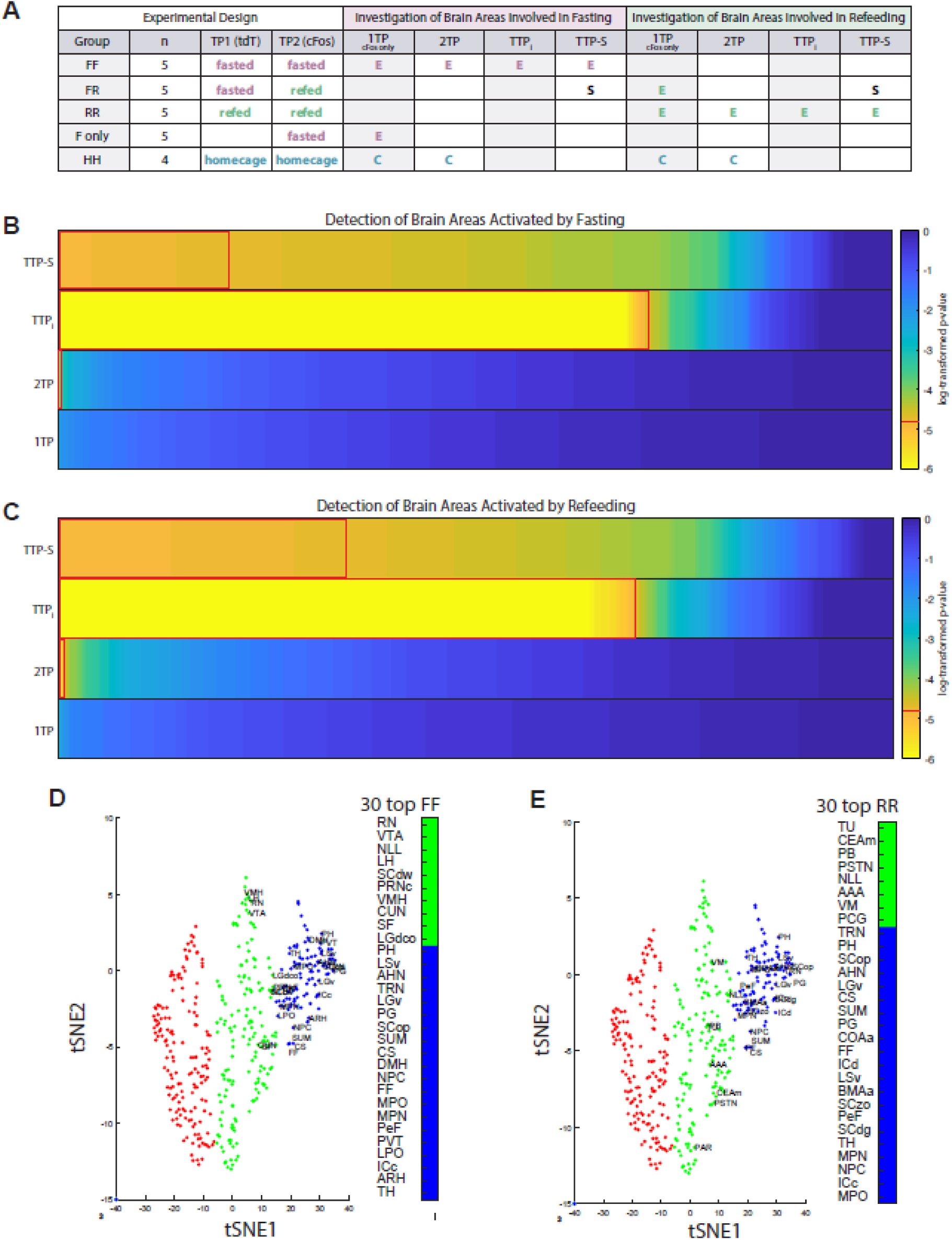
Comparison of methods to identify brain areas activated by fasting vs. refeeding. **A)** Experimental design of one- and two-timepoint assays. Groups of Trap2:Ai14 mice were exposed to stimuli at timepoint 1 (tdTomato) or timepoint 2 (c-Fos). *P*-values derived from different methods were compared for 1TP (using c-Fos), 2TP (using raw counts of double-labeled cells), TTPi (statistical inference) and TTP-S. For each method, the specific groups and numbers of mice used for detecting brain areas activated during fasting or refeeding are indicated in the center and left panels of the grid, respectively. E = experimental group; C = control group; S = subtraction group. 1TP and 2TP comparisons employ E and C groups. TTPi uses only an E group. TTP-S uses an E and an S group. **B)** Comparison of ability of 1TP, 2TP, TTPi and TTP-S to detect brain areas significantly activated during fasting. *P*-values are shown in a heatmap sorted from lowest-to-highest with significant values (*p*<1.6129e-5) outlined in red (also see red line in legend for *p*-value cutoff). **C)** Same as B) for detecting brain areas significantly activated by refeeding. **D**) t-SNE plots showing k-means clustering results of double-labeled cell counts for all mice in all brain areas, with each cluster shown in a different color. The 30 brain areas with the most statistically significant activity in the fasted state as identified by TTP-S are labeled. **E**) Same as D, except the 30 most significantly active brain areas in the refed state are labeled. See Supplemental File S1 for all brain region abbreviations.

### TTP-S Identifies Brain Areas with Intermingled Populations of Differentially Active Neurons in Distinct States

We next sought to compare the patterns of activity across brain areas in the fasted and refed states. These patterns differed substantially (**Figure 5A**), but 36 brain areas were actually activated in both states (**Figure 5B**). A critical question is whether in these 36 brain areas, the fasted and refed states activate the same or different neurons. For example, in the ARH, separate populations of neurons have been shown to be active in each condition (Mandelblat-Cerf et al., 2015). TTP-S was employed to determine if there were fewer double-labeled cells than expected by chance in FR mice, using the HH (homecage-homecage) group of mice for the subtraction. Twenty-six of the 36 brain areas active in both states were identified as having fewer double-labeled cells than expected by chance (72% of brain areas; Fisher’s exact test for overrepresentation between brain areas identified as activated in both states [intersection of FF-FR and RR-FR] and those identified by the FR-HH analysis, *p*=9.4x10^7^; **Figure 5C**). We plotted c-Fos counts of 12 of the 26 brain areas for visualization purposes. Consistent with predictions from our simulations, c-Fos screening alone could not detect significant differences in activity in any of the areas for the fasted and refed states (**Figure 5D**). Further, TTP_i_ identified only 9 out of the 36 brain areas as showing significant lack of double-labeled neurons (Fisher’s exact test *p*=0.09), likely due to its susceptibility to false negatives (**Figure 3B**).

**Figure 5.**
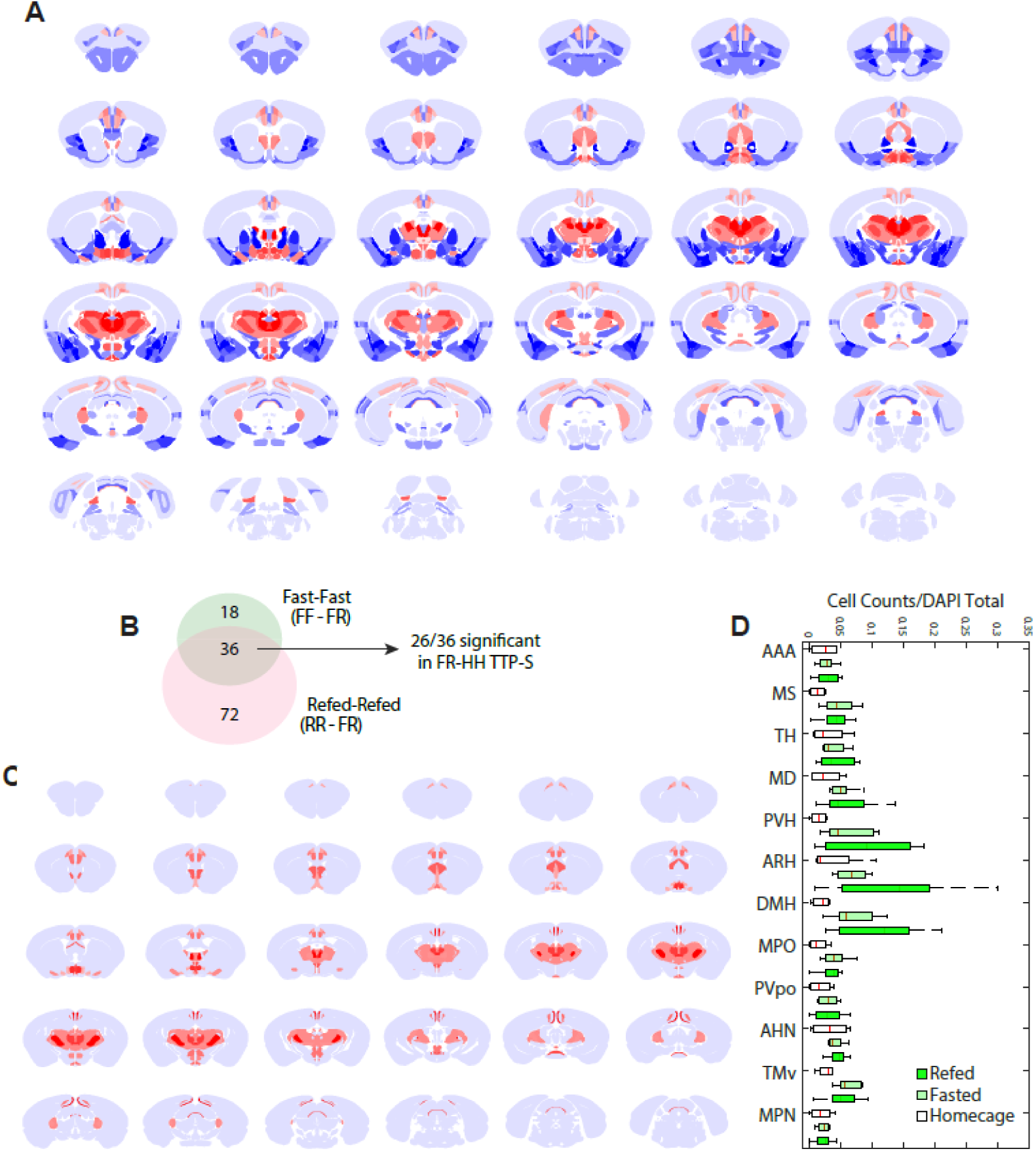
Whole-brain TTP approach identifies areas with intermingled populations of neurons activated neurons during the fasted and refed states. **A)** Allen Brain Atlas heatmap representation of areas significantly activated by refeeding (blue) and fasting (red), as detected by TTP-S (FF-FR, n=5 FF, n=5 FR; RR-FR, n=5 RR, n=5 FR). **B)** Venn diagram showing number of brain areas active in the refed and/or fasted states. 26/36 brain areas active in both states exhibited statistically significant lack of overlap using TTP-S (FR-HH). **C)** ABA heatmap of the 26 brain areas identified as having intermingled populations of neurons activated in the fasted and refed states. **D)** c-Fos counts in fasted, refed, and homecage mice for the 12 most statistically significant brain areas in FF-FR analysis that were also identified as having functionally intermingled populations (n=4-10 mice per group). No brain area had a significant difference in c-Fos counts across conditions (p>0.05, ANOVA or t-test). Brain Areas: **AAA** – Anterior Amygdalar Area; **MS** – Medial Septal Nucleus; **TH** – Thalamus; **MD** – Mediodorsal nucleus of thalamus; **PVH** – Paraventricular hypothalamic nucleus; **ARH** – Arcuate Hypothalamus; **DMH** – Dorsomedial Hypothalamus; **MPO** – Medial Preoptic Nucleus; **PVpo** – Periventricular Hypothalamic Nucleus, preoptic part; **AHN** – Anterior Hypothalamic Nucleus; **TMv** – Tuberomammillary Nucleus, ventral part; **MPN** – Medial Preoptic Nucleus.

### Graph theoretical analysis to identify key circuit nodes exhibiting state-dependent correlated activity

Given that the whole-brain pipeline in combination with TTP-S identified 54 brain areas activated in the hunger state and 108 areas in the refed state, we sought a means to identify which of these identified brain areas are more likely to be key circuit nodes in each state. We therefore turned to graph theoretical analysis (Bassett and Sporns, 2017) to examine data from mice in these two opposing internal states. We applied this analysis to the 126 brain areas activated by either the refed or fasted state (the union of brain areas identified by the FF-FR and RR-FR TTP-S analyses).

The correlation matrices of the number of double-labeled neurons across mice in these 126 brain areas were first examined in both the FF and RR groups of mice. These correlation matrices quantify across mice the degree to which the number of neurons activated at both timepoints in a given brain area are correlated or anti-correlated with the number of neurons activated at both timepoints in other brain areas. Hierarchical clusters of the matrices for the network in the fasted and refed states exhibited different modularity structures (**Figure 6A,B**). Indeed, activity in the same brain areas, - such as the ARH and supramammillary nucleus (SUM) - tended to correlate or anticorrelate with different brain areas in each of the two states (Fisher’s exact test for overrepresentation: ARH *p*=0.17, SUM *p*=1; **Figure 6A,B**), suggestive of differential state-dependent circuit interactions for activated brain areas.

While the correlation matrices indicate that activity in brain areas has state-dependent statistical relationships with activity in other brain areas, it does not itself rank the brain areas in a manner that would enable investigators to preferentially target one or more particular areas for future investigation. To obtain a rank ordering of brain areas to potentially target, we therefore used a PageRank analysis, which was performed separately for data from mice in the fasted and refed states, and separately for the correlation and anti-correlation matrices. PageRank takes into account not just the strength of the correlation between activated neurons in one brain area with another brain area but also the number and strength of correlations across all brain areas, analogous to a website that receives many links to other websites (and has many other websites link to it) and thereby has a high PageRank (see Methods) (Gleich, 2015). **Figure 6C,D** plots the PageRank score for both the correlated and anti-correlated networks from highest score to lowest score. The negative correlation matrices revealed clearly dominant nodes for both the fasted and the refed states, as a subset of brain areas had markedly higher PageRank scores. By contrast, the positive correlation matrices in both states were more similar across all brain areas, and they therefore did not provide a compelling means to target one brain area compared to another (**Figure 6C, D**).

**Figure 6.**
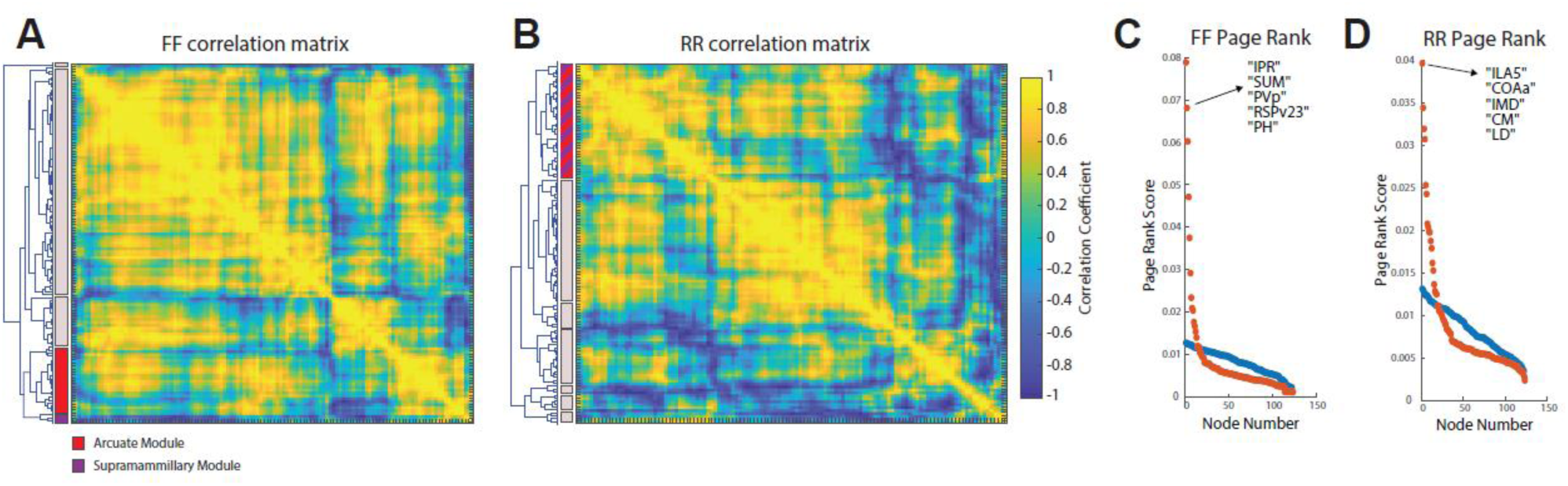
Graph theoretical analysis to identify state-dependent brain regions acting as key network nodes. **A)** Correlation matrix of double-labeled counts in fast-fast (FF) mice (n=5) in the 126 brain areas that were identified as significant in both the FF-FR and RR-FR analyses. Hierarchical clustering determined the location of brain areas on the matrix (see clustering on y-axis) and modules are noted by the bars on the y-axis. Red indicates the module containing the arcuate nucleus of the hypothalamus (ARH), and purple indicates the module containing the supramammillary nucleus (SUM). **B**) Same as A), but for RR mice (n=5). Note that the ARH and SUM are now in the same module, indicating that the correlation structure of activity across brain areas differs in the fasted vs. refed states. **C**) PageRank analysis (see Methods) to show the importance of brain regions ranked according to positive (blue dots) and negative (red dots) correlations with other brain regions in FF mice. The top 5 most important brain areas by PageRank (all from the negative correlation network) are labeled. **D**) Same as C) but for RR mice. Brain Areas: **IPR** - Interpeduncular nucleus, rostral; **SUM** – supramammillary nucleus; **PVp** – Periventricular hypothalamic nucleus, posterior part; **RSPv23** - Retrosplenial area, ventral part, layer 2/3; **PH** – Posterior hypothalamic nucleus**; ILA5** – infralimbic area, layer 5; **COAa** – cortical amygdalar area; **IMD** - Intermediodorsal nucleus of the thalamus; **CM** - Central medial nucleus of the thalamus; **LD** - Lateral dorsal nucleus of thalamus.

The PageRank score computed by applying graph theoretical analysis provides an unbiased means of rank ordering brain areas in a manner that could guide future investigations. However, when an experiment is designed, as the current one was, to study two opposing states likely to engage neural circuits in different ways, it is also important to establish that a brain area exhibits state-dependent properties. To explore the extent to which this was true in our dataset, we plotted the network topologies of both the positive and negative correlation matrices in the fasted and refed states (**Figure 7**). In these plots, each brain area is plotted with a circle whose diameter is proportional to its PageRank. Further, in **Figure 7A-D**, we highlight the SUM and brain areas correlated with activity in this area. The SUM was shown to have more double-labeled cells than expected by chance in both the fasted and refed states, and it was shown to have fewer double-labeled cells in FR mice than expected by chance. Further, the SUM nucleus was one of the brain areas with the highest PageRank score in the negative correlation matrix in the fasted state. Activity in SUM was anticorrelated with a large number of brain areas during the fasted state, and many of these brain areas were brain areas identified as being active in the refed state (**Figure 7A-D**). Thus, the network topology demonstrates state-dependence with respect to which brain areas are correlated or anticorrelated with activity in the SUM. Note that state-dependent network topologies can often be observed in many brain areas, particularly those with distinct populations of neurons active in the fasted and refed states. For example, examination of network topologies with the ARH highlighted, along with brain areas correlated or anticorrelated with the ARH, also demonstrates clear state-dependence (**Figure S5**).

**Figure 7.**
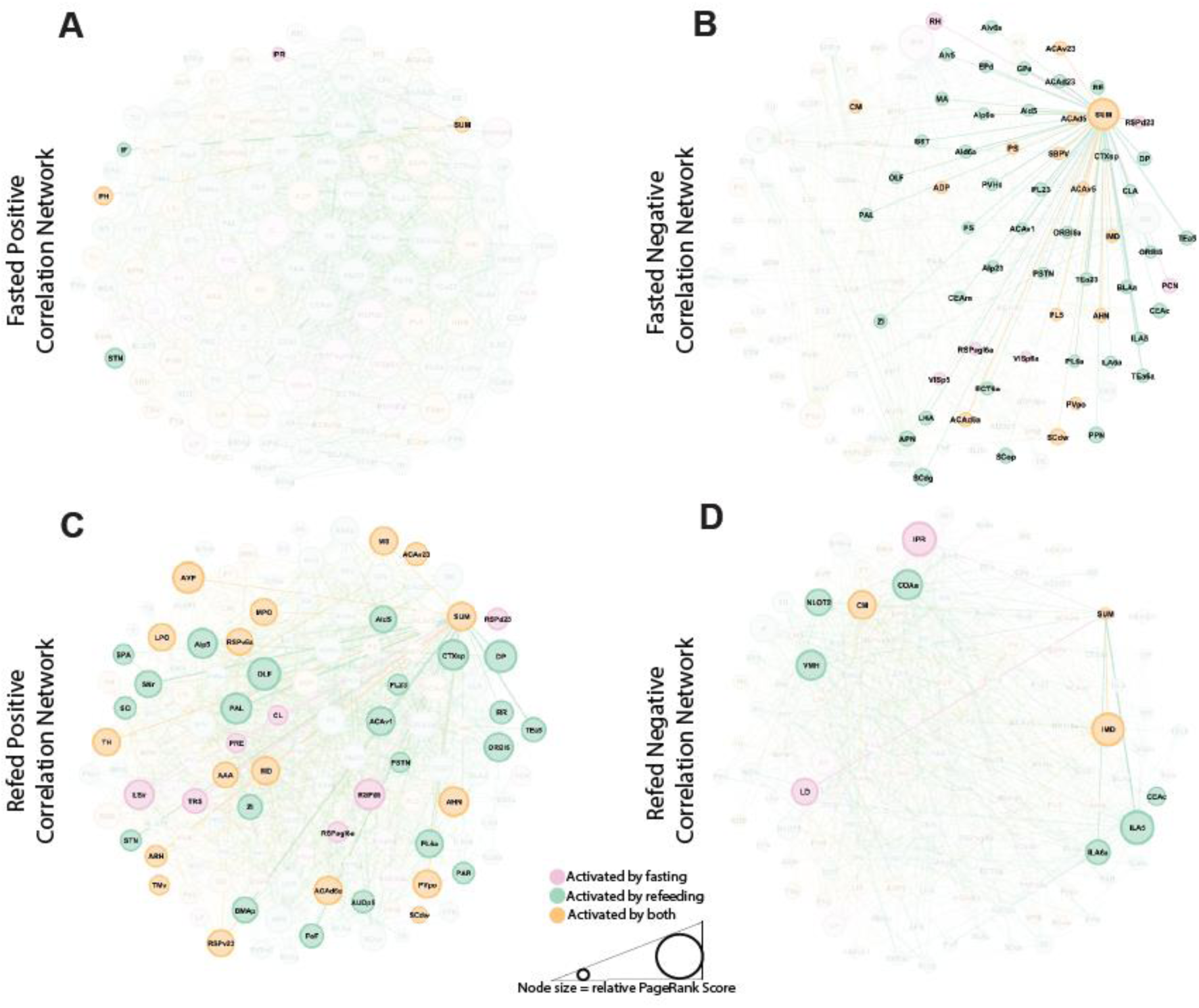
Graph theoretical analysis to visualize functional networks activated during fasting and refeeding. Graph theoretical approaches were used to visualize the network topology of neurons activated by either the fasted or refed states (126 brain areas). **A)** Positively-correlated network connectivity for the supramammillary nucleus (SUM) in the fasted state (FF mice; n=5). Each line indicates a positive correlation (R>0.6) between brain areas [Fruchterman-Reingold layout]; Node colors indicate whether brain areas were significantly activated by fasting (red/pink), refeeding (green) or both (orange). Only nodes positively correlated with SUM are highlighted. The size of the node indicates the PageRank score, an indication of the connectivity of each node to the network. **B**) Negatively-correlated network connectivity of SUM in the Fasted state (FF mice; n=5). Each line indicates anticorrelated activity (R<-0.6) between brain areas. **C**) Same as E) for SUM except in refed state. **D**) Same as F) for SUM except in refed state. See Supplemental Information for all other brain region abbreviations.

### Whole-brain and graph theoretical approaches to identify circuits activated by semaglutide

Satiety and hunger can be induced by pharmacologic means, though the brain circuits through which drugs like the GLP1 agonist semaglutide work to reduce food intake remain an active area of investigation (Gabery et al., 2020; Kabahizi et al., 2022). As a first step towards applying the methodology described in this paper to identify brain areas and networks activated by semaglutide, we performed our whole-brain pipeline along with TTP-S in groups of mice where semaglutide (10nmol/kg) was administered at each of two timepoints. For these experiments, mice for the subtraction group were simply in the sated condition while in the homecage at both time points (HH mice shown in **Figure 4A**). Results were compared to mice administered the hormone ghrelin (10nmol), known to induce hunger in humans and rodents (Egecioglu et al., 2010; Levin et al., 2006; Tschöp et al., 2000).

TTP-S identified 24 areas activated by semaglutide within one hour of treatment (**Figure 8A,C**). These 24 areas were exclusive of the hindbrain and SFO, which for technical reasons already described were not included in analyses. By comparison, a four-hour acute treatment with semaglutide was previously reported to activate 5 areas (excluding the hindbrain and the SFO) using standard 1TP to analyze whole brains (Gabery et al., 2020). TTP-S analysis identified four of these five areas (bed nucleus of stria terminalis, central amygdala, parasubthalamic nucleus & midline thalamic nucleus). The vascular organ of laminar terminalis was the one area activated by four-hour treatment in (Gabery et al., 2020) but not in a one-hour treatment according to our analysis. However, immediately adjacent areas were activated. Overall, there was statistically significant overrepresentation in the brain areas identified as being activated by semaglutide in our study and a prior study surveying the whole brain for activation by semaglutide (assuming a screen of 570 brain areas in both studies, *p*=5.0x10^-7^, Fisher’s exact test).

Strikingly, we observed that a large and statistically significant fraction of the brain areas activated by semaglutide were also activated during refeeding (18/24 brain areas (75%), Fisher’s exact test *p*=1.4x10^-9^, **Figure 8A**). These data suggest that semaglutide administration may induce a brain state that shares features of the state induced by refeeding (satiety). The similarity between states induced by refeeding and by semaglutide delivery could also be appreciated by examining c-Fos expression and counts of double-labeled cells across mice in brain areas commonly activated by the two states (**Figure S6A,B**). By contrast, a large majority of the 35 areas activated by ghrelin did not overlap with those areas activated in the fasted state (4/35 brain areas (11.4%), Fisher’s exact test *p*=0.43, **Figure 8B**). This raises the possibility that, at least at short-time scales in relation to activity induced by ghrelin administration, the brain state associated with hunger is not mimicked. Nonetheless, there was little overlap between areas activated by semaglutide and ghrelin (**Figure 8C**). One can also visualize the difference in activity induced by semaglutide and ghrelin in these brain areas by examining double-labeled cell counts (**Figure S6B**).

We next sought to use graph theoretical analysis to rank order the brain areas activated by semaglutide (24 brain areas) and ghrelin (35 brain areas). As in the fasted and refed states, the negative correlation matrices proved to be the most revealing in terms of identifying areas whose activity was most tightly correlated or anti-correlated with other brain areas across mice. One striking finding was that in the semaglutide condition, the lateral subdivision of the central nucleus of the amygdala (CEAl) had a high PageRank score in the negative correlation matrix (**Figure 8D,E**). In the ghrelin condition, ARH stood out as having a high PageRank score in the negative correlation matrix (**Figure 8G,H**). **Figure 8E,H** shows the negatively-correlated network topologies for the ghrelin and semaglutide groups of mice, highlighting the relationship between CEA1 and ARH and other brain areas whose activity is statistically related to each of the two areas. For comparison, we also depict network topologies in the homecage condition for the same brain areas (**Figure 8F,I**), as well as the positively correlated networks (**Figure S7A-D**). These data reveal that the networks correlated with CEA1 and ARH are state-dependent. Overall, the TTP-S whole-brain pipeline, combined with graph theoretical analysis using PageRank scores, identifies candidate brain areas for future investigation based not only on a brain area’s activity in relation to a state or event. Instead, the approach takes into account correlation matrices describing the relationship between activity across brain areas, and it takes into account whether a brain area exhibits state-dependent network topology.

**Figure 8.**
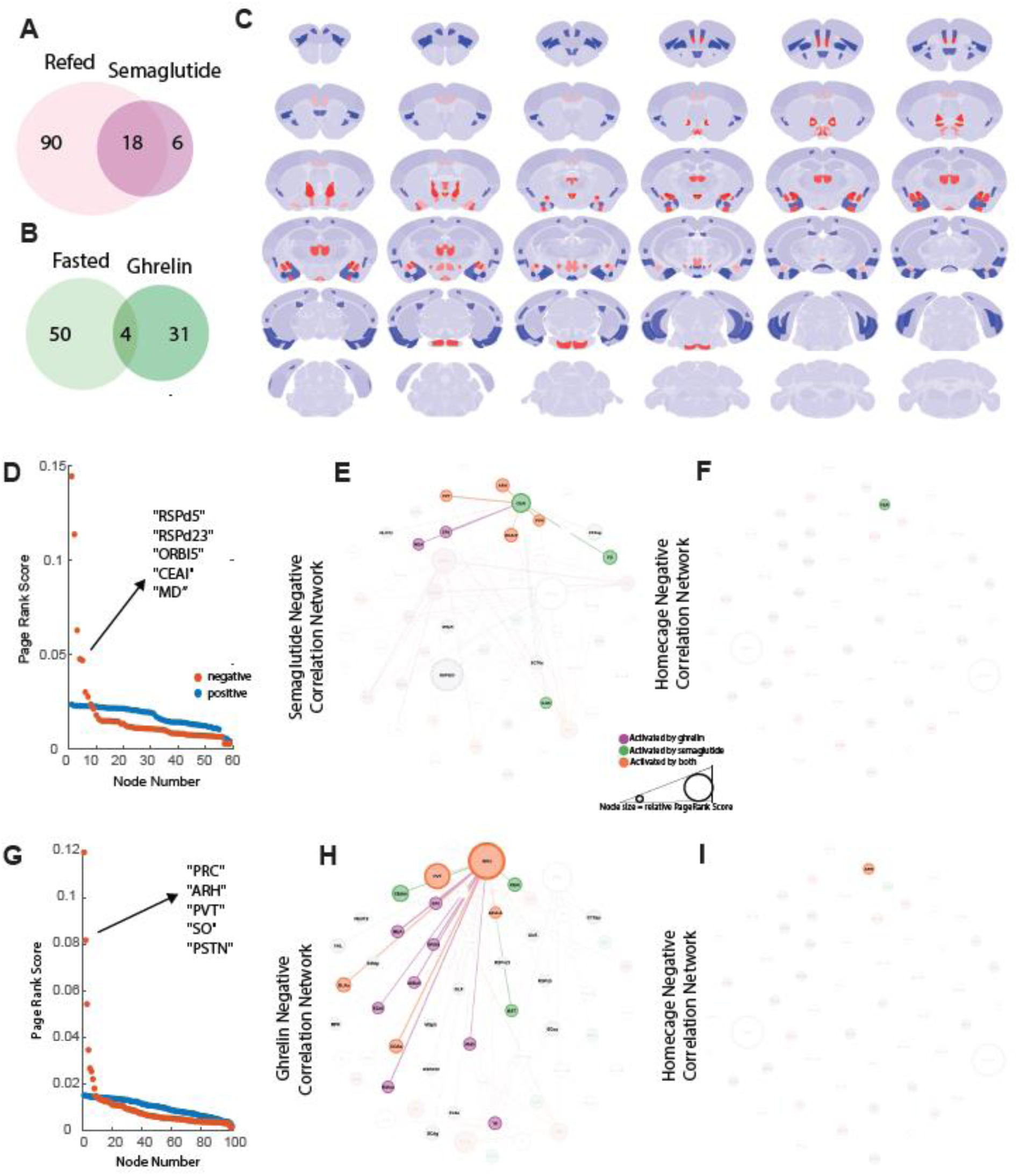
Graph theoretical analysis to visualize functional networks activated semaglutide and ghrelin. **A**) Venn diagram showing number of brain areas significantly activated by the refed (TTP-S using RR-RF mice) and semaglutide administration (TTP-S using SS-HH mice) states (n=4-5 in each group). **B**) Venn diagram showing number of brain areas significantly activated by the refed (TTP-S using FF-RF mice) and ghrelin administration (TTP-S using GG-HH mice) states (n=4-5 in each group). **C**) Allen Brain Atlas heatmap representation of areas significantly activated by semaglutide (red) and ghrelin (red) in the TTP-S analyses for each group (SS-HH and GG-HH). **D**) PageRank analysis of data from negative (red) and positive (blue) correlation matrices in SS mice (n=4). **E**) Negatively-correlated network connectivity for the lateral subdivision of the central amygdala (CEAl) in SS mice (n=4). Each line indicates anticorrelated activity (R<-0.6) between brain areas [Fruchterman-Reingold layout]; Node colors indicate whether brain areas were significantly activated by semaglutide (green), ghrelin (purple), or both (orange). Only nodes negatively correlated with CEAl are highlighted. The size of the node indicates the PageRank score, an indication of the connectivity of each node to the network. **F**) Negatively-correlated network connectivity (or lack thereof) for the CEAl in HH control mice (n=4). **G**) PageRank analysis of data from negative (red) and positive (blue) correlation matrices in ghrelin-ghrelin exposed mice (n=5). **H**) Negatively-correlated network connectivity for the arcuate nucleus of the hypothalamus (ARH) in GG mice (n=5). Each line indicates anticorrelated activity (R<-0.6) between brain areas. Only nodes negatively correlated with ARH are highlighted. The size of the node indicates the PageRank score, an indication of the connectivity of each node to the network. **I**) Negatively-correlated network connectivity (or lack thereof) for the ARH in HH control mice (n=4). Brain Areas: **RSPd5** - Retrosplenial area, dorsal part, layer 5; **RSPd23** - Retrosplenial area, dorsal part, layer 2/3; **ORBl5** - Orbital area, lateral part, layer 5; **CEAl** - Central amygdalar nucleus, lateral part; **MD** - Mediodorsal nucleus of thalamus; **PRC** - Precommissural nucleus; **ARH** - Arcuate hypothalamic nucleus; **PVT** - Paraventricular nucleus of the thalamus; **SO** - Supraoptic nucleus; **PSTN** - Parasubthalamic nucleus. See Supplemental Information for all brain region abbreviations.

## Discussion

Advances in neurotechnology have made it possible to monitor and manipulate the activity of hundreds to thousands of neurons simultaneously in multiple targeted brain areas in order to elucidate mechanisms underlying different cognitive, emotional, physiological and behavioral processes. This enhanced capability makes it more important than ever to develop approaches for determining which brain areas to target for study with these new technologies. Current approaches for whole-brain screening typically rely on IEG expression at a single timepoint, which requires prohibitively large sample sizes to detect significant activity after multiple comparisons correction. Here we present a novel methodology for screening the whole brain for areas where single neurons are activated by particular stimuli or events. Our approach combines a number of critical elements: 1) the use of a transgenic mouse line (TRAP2 mice) that enables labeling of activated neurons at two timepoints, not one; 2) a pipeline that uses serial sections across the whole brain, software that performs registration to the Allen Brain Atlas Common Coordinate Framework, and deep learning-based image restoration and cell detection; 3) statistical analyses that assess whether a brain area exhibits more or fewer cells activated at each of two timepoints than expected by chance; 4) strategic experimental designs to increase the sensitivity and specificity of screening by utilizing and comparing two groups of mice who experience different stimuli at the two timepoints; and 5) the application of graph theoretical analysis to rank order brain areas according to the degree of correlated or anticorrelated activation with other identified brain areas. We believe this approach will have wide applicability for systems neuroscience research that seeks to prioritize brain areas for study in understanding neural circuit function in relation to a wide variety of states and processes.

In this paper, we validate the novel whole-brain approach by studying neural networks activated by two opposing states: hunger and satiation. During periods of fasting, decreases in circulating hormonal factors including leptin and insulin produce hunger in animals, with a concomitant coordinated response in the body to obtain calories and nutrients (Morton et al., 2006; Morton et al., 2014; Spiegelman and Flier, 2001). Prior research has shown that this coordinated response engages neurons in the hypothalamus involved in motivation and appetite (Caron and Richard, 2017; Rossi and Stuber, 2018; Sternson and Eiselt, 2017), areas of the motor cortex engaged in foraging behaviors (Dietrich et al., 2015; Watts et al., 2022), and sensory cortical areas involved in detecting food and food cues (Burgess et al., 2016; Livneh et al., 2017), among other brain areas. Conversely, satiety is accompanied by an increase in bodily hormones that signal fullness to the brain (Beutler et al., 2017; Sternson and Eiselt, 2017), engaging areas of the brainstem and midbrain (Carter et al., 2013; Chaudhri et al., 2008; Kim et al., 2020), as well as corresponding motor and sensory responses involved in rest and digestion (Watts et al., 2022). Thus, a large number of brain areas are known to be active during both internal states, hunger and satiety, suggesting that these states could serve well to validate our methods.

Despite using a small sample size of 5 mice in a group, with statistical assessments typically involving comparison of two groups, the whole-brain pipeline we describe identified a large number of the brain areas previously implicated in satiation, hunger and feeding. Traditional screening methods using activity-dependent expression of IEGs at a single timepoint using these sample sizes would not have identified any of these brain areas upon correction for multiple comparisons. Further, the approach developed not only identified brain areas activated by hunger and satiety, but it also determined if a brain area possessed distinct populations of neurons activated by each of the two states. The paradigmatic demonstration of this ability was identification of the ARH, an area where prior work has demonstrated intermingled circuits activated in the hunger and sated states, respectively (Mandelblat-Cerf et al., 2015). Single timepoint methods for whole-brain screening for activated neurons cannot identify these types of intermingled circuits, as they cannot determine if the same or different neurons are activated by two different stimuli or events. Moreover, methods of assaying brain activity at a coarser spatial scale than the single cell, such as with fMRI, also cannot determine if the same or different cells are being activated by two distinct events.

Our approach differs from several recent tools that allow for high throughput imaging and analysis across a whole rodent brain (Ragan et al., 2012; Seiriki et al., 2017; Stelzer, 2015; Ueda et al., 2020). The most commonly used method combines labeling of IEGs such as c-Fos with whole brain clearing and imaging to detect activated cells (e.g., iDISCO/iDISCO+ (Renier et al., 2016; Renier et al., 2014)). The main advantage of these other approaches is the ability to keep brain tissue intact, which facilitates accurate registration to a common coordinate framework. However, inconsistent antibody penetration exacerbates variability in cell counts both across and within brains (Molbay et al., 2021; Terstege and Epp, 2022). Moreover, obtaining accurate total cell counts within each brain area typically requires imaging at resolutions that demand prohibitively long acquisition times on commonly used light-sheet microscopes, presenting a major obstacle to screening or network analysis. However, recent developments in light-sheet microscopy, enabling higher resolution and faster acquisition speeds, may render this technique more broadly feasible for intact whole-brain studies.

The whole-brain screening described here uses traditional serial sections and confocal or widefield fluorescence microscopy, a more accessible technique which minimizes challenges in immunolabeling and provides accurate cell counts. These cell counts include the number of labeled cells at each of two timepoints, and the total number of cells in each brain area within individual animals. By obtaining these cell counts, we were able to implement statistical approaches to determine if a particular stimulus or event activates neurons in a brain area using relatively small sample sizes. However, and unfortunately, in its current iteration occasional errors in the registration process occur in some parts of the brain that tend to be distorted or damaged. As a result, poorly registered brain areas were eliminated from our analyses, and these excluded areas typically included the cerebellum and much of the hindbrain. Future experiments could employ refined versions of clearing protocols such as iDISCO+, EZ Clear, and WildDISCO (Hsu et al., 2022; Mai et al., 2024; Renier et al., 2016; Renier et al., 2014), or commercial methods like LifeCanvas (Tian et al., 2023), to provide consistent and homogenous labeling of all neurons, in addition to c-Fos, within intact brains. Coupled with newly available light-sheet microscopes, the statistical approaches used here will be an even more powerful tool for efficient and highly sensitive whole-brain screening.

Just like for hunger and satiety in this study, many cognitive, emotional and behavioral processes almost certainly also engage large numbers of brain areas. The number of activated brain areas identified demands an approach for prioritizing for future study those areas among the activated brain areas. To accomplish this prioritization, we applied graph theoretical analysis. This analysis was applied to data in which neurons were activated at each of two timepoints in relation to satiety or hunger. Graph theoretical analysis provides a means for rank ordering the activated brain areas according to the degree to which activity (cells active at two timepoints in our case) in each brain area is correlated (or anti-correlated) with activity in other brain areas across mice (the PageRank score) (Bassett and Sporns, 2017; Gleich, 2015). In the context of hunger and satiety, the top-ranked brain areas identified by graph theoretical analysis exhibited state-dependent correlations with distinct groups of brain areas, many of which were known to be involved in these processes according to prior studies (Andermann and Lowell, 2017; Mandelblat-Cerf et al., 2015; Morton et al., 2014; Myers and Olson, 2012; Sternson and Eiselt, 2017; Watts et al., 2022). We propose that statistically significant activation, a high PageRank score using graph theoretical analysis, and a state-dependent statistical relationship with activity in distinct networks of brain areas (distinct modules) constitute criteria for prioritizing a brain area for future study. Together, this approach promises to more completely identify the complex neural circuits that operate at the brain-wide level in relation to many behaviors and internal states.

To explore another potential application of our methods, we examined how semaglutide administration increases activity in the brain. Semaglutide and related GLP1 agonists are transforming the treatment of obesity (Lincoff et al., 2023; Wilding et al., 2021), yet a mechanistic understanding of how these new drugs impact feeding circuitry remains incomplete. Some studies have screened for activity using single timepoint activity in relation to semaglutide administration, identifying a range of brain areas as candidate structures (Gabery et al., 2020). Some of these brain areas are in the hindbrain, a part of the brain where the current iteration of our whole-brain pipeline does not survey due to damage and distortion and concomitant difficulty with registration to the Allen Brain Atlas. Nonetheless, many of the other brain areas identified previously were also identified with our whole-brain pipeline. Further, the brain areas activated by semaglutide delivery were statistically associated with those areas activated in the sated state. Finally, the application of graph theoretical analysis suggested that the CEAl might be a prime circuit node mediating semaglutide’s efficacy, due its high PageRank score that was also state-dependent. This discovery contrasted with the finding that ARH is an important node upon ghrelin administration, a hormone generally considered to induce hunger.

Despite the power of the whole-brain screening pipeline and associated analyses described here, the approach suffers from some limitations. First, our approach, like all methods based on IEG expression, is limited by the faithfulness (or lack thereof) of IEG expression in reflecting neural activity. Second, the use of molecular labels of neural activity based on IEG expression lacks temporal specificity, so activity over a relatively long timescale might result in labeled cells. This means cells might be activated and labeled by many different stimuli or events despite best efforts otherwise. New methods for molecular labeling of active cells are emerging (Shi et al., 2024), but at this stage these approaches are not suitable for application in whole-brain screening. Third, some states or events may cause neurons to decrease their firing rate. This need not preclude the use of our whole-brain pipeline, but the application might require larger sample sizes. For example, if the hunger state decreases activity in a brain area, the number of cells activated in the hunger state might be fewer than the number activated in that area when mice are not hungry. With a sufficient sample size, the whole-brain approach described here could detect this difference, so long as the decrease in activity was sufficient to eliminate activity dependent expression of IEGs in some cells. This type of decrease in activity could account for how graph theoretical analysis identifies brain areas where higher activity in one brain area is anti-correlated with activity in other brain areas (high PageRank score for negative correlation matrices).

Another potential limitation of the whole-brain method is that some states or events may modulate neural activity only in particular conditions without changing neurons’ activity in other conditions. This type of modulation has been reported with respect to semaglutide delivery, where activity in neurons in the DMH of the hypothalamus is modulated only when mice are engaged in behavior related to food (Kim et al., 2024). By contrast, semaglutide does not modulate spontaneous DMH neuronal activity. In principle, since this type of modulation involves an increase in activity, the whole-brain pipeline could detect brain areas where the modulation of activity is condition-dependent, as in the case of semaglutide in DMH. This will particularly be true if an increase in activity requires two conditions to be met (e.g. if increases in activity in some DMH neurons require a food-related condition and semaglutide delivery). In this case, experimental designs that include groups of mice where both conditions are met, as well as groups where both conditions are not met, could be used to exploit the whole-brain pipeline to identify brain areas where activity is modulated.

In human neuroscience, whole-brain neuroimaging has been the backbone of research since the development of fMRI (Raichle, 2009). Despite the power of fMRI studies, some researchers have emphasized challenges with respect to the reproducibility of findings, to the prevalence of small effect sizes, and to the difficulty in identifying clinically significant diagnostic markers (Bennett et al., 2009; Etkin, 2020; Marek et al., 2022). These challenges in part may be due to the lack of cellular level resolution when using fMRI. However, robust whole-brain screening methods in animal models have suffered from their own challenges, including issues related to required sample sizes. Our approach for whole-brain screening proposes specific criteria for identifying important brain areas for a particular function. First, a brain area should be activated by a particular state or event. Second, brain areas that exhibit high statistical correlation or anti-correlation with activity in other brain structures might be particularly important to target. Third, the statistical relationship with activity in other brain areas should be state-dependent, thereby conferring functional specificity to the finding. Together these criteria promise to help identify key neural circuits and nodes across a broad range of internal states and behaviors, which could be key for understanding mechanisms underlying therapeutic interventions for many different disorders.

## Methods

### Experimental Animals

Homozygous Fos2A-iCreER/2A-iCreER (TRAP2) mice (Allen et al., 2019) were obtained from Jackson Labs (stock #030323), and R26Ai14/+ (Ai14) mice (Madisen et al., 2010) were obtained from Jackson Labs (stock #007914). TRAP2 mice were crossed to Ai14 mice to obtain the double heterozygous (TRAP2:Ai14) that were used for all TTP experiments. Littermate mice of both biological sexes were used in all experiments and were between 8-12 weeks of age. Mice were single-housed during all experimental procedures. All procedures were performed in accordance with standard ethical guidelines approved by Columbia University Institutional Animal Care and Use Committee.

### Drug Preparation

**4-hydroxytamoxifen** (4-OHT; Sigma, #H6278) was dissolved in ethanol at a concentration of 20mg/ml and then added to a 1:4 mixture of castor oil:sunflower seed oil (Sigma, #259853 and #S5007), and the ethanol was evaporated by vacuum centrifugation to arrive at a final concentration of 10mg/ml 4-OHT. 4-OHT solutions were always prepared on the day of use. Mice were injected intraperitoneally (i.p.) at a concentration of 60mg/kg, similar to previous studies using TRAP2 transgenic mice (Allen et al., 2019; DeNardo et al., 2019). **Ghrelin (**Bachem #4033076.0500) was dissolved in sterile saline to a concentration of 1ug/ul (equivalent to 0.3nmol/ul) and aliquoted in 50ul tubes for storage at -20°C. On the day of use, aliquots were thawed, and each mouse was injected i.p. with 10nmol of drug, similar to previous studies (Andrews et al., 2009; Wren et al., 2001). **Semaglutide** (Bachem, #4091661) was dissolved in 5% mannitol for a stock solution of 20nmol/ml and stored at -20°C in 200ul aliquots. On the day of use, 200ul aliquots were diluted into 1.8ml of mannitol solution to make a concentration of 2nmol/ml. Animals were injected subcutaneously with 10nmol/kg, similar to previous studies (Gabery et al., 2020).

### Two-Timepoint Behavioral Assays

8–12-week-old heterozygous TRAP2:Ai14 animals were single-housed and habituated to a reverse-light cycle room (12hr:12hr) for at least 1 week. For the **Fast-Refed Two-Timepoint Experiments,** mice were split into 4 groups of 5, age and weight matched across groups. Mice were food restricted for 18 hours prior to being injected with 4-OHT (for timepoint 1 of our TTP assays) at the beginning of their dark-cycle. Food was only re-introduced 8 hours after 4-OHT injections, whereafter we resumed an *ad-libitum* food schedule. All mice were habituated to i.p. injections with sham saline injections for at least 5 days prior to being given 4-OHT at timepoint 1. Four weeks later, the mice were fasted for 24 hours, then refed for 2 hours (in the case of the Fast-Refed or FR group; n=5) prior to being sacrificed for brain harvesting and histology. The amount of food consumed was measured by weighing pellets before and after refeeding. The same protocol was followed for the Refed-Fast (FR; n=5) assays except the order was reversed, and similarly for the Fast-Fast (FF; n=5) and Refed-Refed (RR; n=5) conditions. Homecage-Homecage controls (HH; n=4) were performed where mice were not fasted or refed and always had continuous access to *ad-libitum* food access. These mice were similarly habituated to i.p. injections for at least 5 days prior to receiving 4-OHT in a homecage condition at timepoint 1. They were then allowed to *ad-libitum* access to food for another 4 weeks prior to sacrifice and brain harvesting for histology.

### Ghrelin TTP Assays

Age- and weight-matched mice were housed in a reverse-light cycle room, and food intake and body weight were measured daily. Mice were given *ad-libitum* access to food and were habituated to i.p. injections with sham saline injections for at least 5 days prior to timepoint 1. For the first timepoint, food hoppers were removed from the cage ∼3hrs before mice received an i.p. injection of 10nmol ghrelin followed immediately after by 60mg/kg 4-OHT given i.p. Food hoppers were placed back into the cages ∼8hrs after injections – this was done to ensure that ensembles related to eating would not be captured in our whole-brain screening. 2 weeks later, for timepoint 2, mice received another i.p. injection of 10nmol ghrelin and were sacrificed for c-Fos staining 1.5hrs later. (Ghrelin-Ghrelin or GG group; n=5).

### Semaglutide TTP Assays

Age- and weight-matched mice (n=4) were housed in a reverse-light cycle room, and food intake and body weight were measured daily. Mice were given *ad-libitum* access to food and were habituated to i.p. injections with sham saline injections for at least 5 days prior to timepoint 1. For the first timepoint, food hoppers were removed from the cage ∼3hrs before mice received a subcutaneous injection of 10nmol/kg semaglutide given simultaneously with 60mg/kg 4-OHT delivered i.p. Food hopper were placed back into the cages ∼8hrs after injections. 2 weeks later, for timepoint 2, mice were injected subcutaneously with 10nmol/kg semaglutide and sacrificed 1hr later for brain harvesting and histology (Semaglutide-Semaglutide or SS group; n=4).

### Histology and Imaging

Mice were transcardially perfused with 4% paraformaldehyde (PFA) and brains were stored in 4% PFA overnight at 4° C. Brains were transferred to PBS for 24 hrs, followed by 20% sucrose in PBS for 48hrs. Brains were then dried and embedded in Tissue-Tek Optimum Cutting Temperature (OCT; Sakura Finetek, #4583) medium prior to flash freezing and storage at - 80°C until sectioning. Brains were sectioned on a cryostat (Leica Microsystems) at 50 microns through the entire brain, and floating sections were collected in PBS. For immunostaining, sections were washed 3 times in PBS and then incubated in 5% donkey serum (Jackson Immunoresearch; #017-000-121) in PBS with Triton-X (PBST) for 1hr. Sections were then stained for using primary antibodies targeting c-Fos (Cell Signaling, #2250, 1:1000 dilution in PBST and 5% donkey serum) and NeuN (EMD Millipore, #MAB377, 1:1000 dilution) and incubated at 4°C overnight. Sections were then washed 3 times with PBST prior to conjugation with secondaries AlexaFluor-647 (Invitrogen #A31573, 1:1000 dilution targeted to anti-c-Fos) and AlexaFluor-488 (Life Technologies #A21202, 1:1000 dilution), as well as a DAPI counterstain (Sigma-Aldrich, #D9542 1:5000). Sections were mounting onto Superfrost Plus slides (Fisher Scientific, #12-550-15) and coverslipped with mounting medium. Confocal images of whole sections were then obtained using a Yokogawa W1 spinning disk confocal microscope using a 4x 0.2 NA objective (for whole-brain analysis) and a 10x 0.45 NA objective (for verification in the arcuate nucleus of the hypothalamus (ARH) shown in Figure 1) tiled to cover the whole section. Images were captured in 4 channels: 405nm (corresponding to DAPI stain), 488nm (corresponding to anti-NeuN stain), 561nm (corresponding to tdTomato (tdT) label induced in TRAP2:Ai14 mice by 4-OHT at timepoint 1), and 647nm (corresponding to anti-c-Fos stain performed at timepoint 2).

### 10x Two-Timepoint Analysis in Arcuate Nucleus

Images from the ARH in FF, FR, RF, and HH mice were analyzed in FIJI (Schindelin et al., 2012). ARH images from the 488nm, 560nm, and 647nm channels (corresponding to anti-NeuN, tdT, and anti-c-Fos, respectively) were manually quantified and examined for double-labeled cells.

### Whole-Brain Analysis Pipeline

Images corresponding to single 50 micron sections through the entire were pre-processed in FIJI/ImageJ in order to renumber the sections sequentially from anterior to posterior and remove background to be compatible with the BrainJ whole-brain registration plugin, which has been described previously (Botta et al., 2020). Briefly, images are centered and rotated to facilitate registration for each individual brain section, which yields a 3D brain volume rendering. Images are then registered to the Allen Brain Atlas Common Coordinate Framework (CCF) using Elastix (Klein et al., 2009) by utilizing the DAPI-stained channel to register each section. Note that we attempted to use NeuN as a marker of total cell count, but were not consistently able to attain accurate counts in an automated manner particularly in brain areas with high neuronal density. Therefore, we utilized the DAPI channel for total cell counts in our whole-brain pipeline. To extend BrainJ’s capabilities for image restoration, deep-learning segmentation and provide additional functions required for this analysis we developed an updated version of BrainJ in Python (Botta et al., 2020) This version of BrainJ automatically detected cells in each channel, and we considered cells double-labeled using an algorithm that combines information on the distance between centroids and fraction of overlapping volume between two cells (i.e. - masks), with thresholds set for each brain region separately. BrainJ generates tubular data for detected cells in all 4 channels in each brain area. The BrainJ plugin for Fiji and information on the updated pipeline for Python can be found at https://github.com/lahammond/BrainJ.

### Statistical methods

#### Two-timepoint Statistical Inference (TTP_i_)

TTP_i_ was performed on individual brains and across registered brain areas by initially assuming a hypergeometric distribution, where the probability distribution is described by:

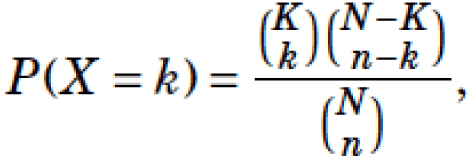

where *X* denotes the random variable giving the size of the overlap, *N* is the total cell count, *K* is the number of active cells at timepoint 1 (TP1; marked by tdT in our *in vivo* whole-brain screens), *n* is the number of labeled cells at timepoint 2 (TP2; marked by c-Fos in our whole-brain screens), and *k* double-labeled cells. The 1-sided *p*-value is given by the probability that *at least* k neurons are double-labeled, or the probability that fewer than k neurons are double-labeled, and is given by the sum of all probabilities either equal to or greater than k, or equal to or less than k:

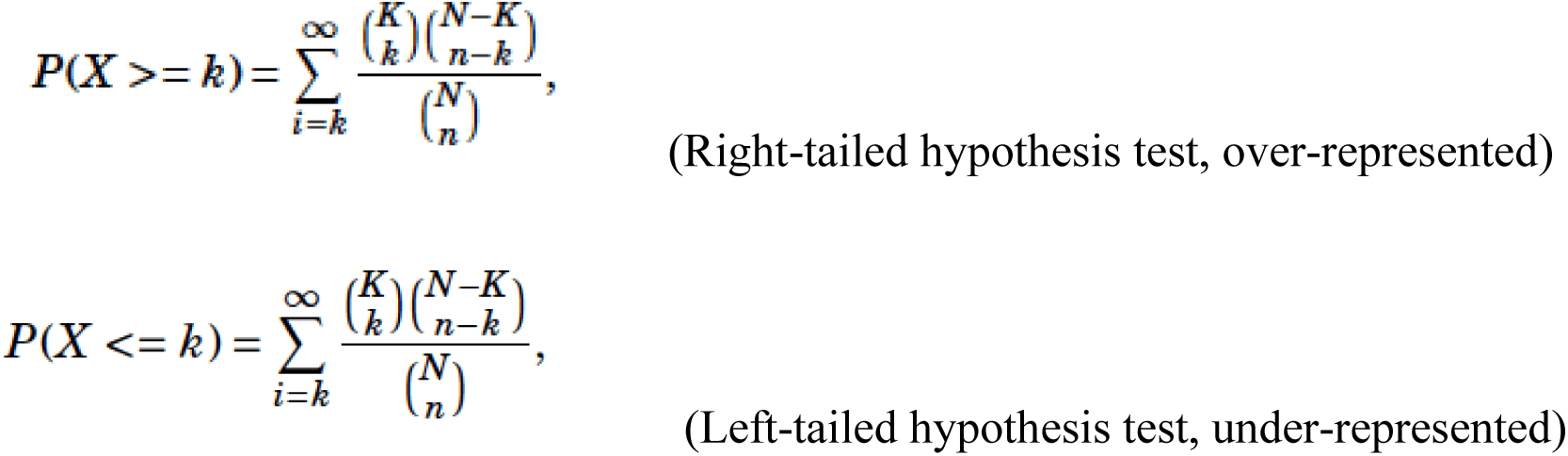

We exploit the equivalence of the 1-sided *p*-value for a hypergeometric distribution and the 1-tailed Fisher’s exact test, which can be implemented in Matlab using a contingency table. For combining *p*-values across animals, we used a harmonic mean approach.

#### Two-timepoint with Subtraction (TTP-S)

TTP_i_ (i.e., without subtraction) was carried out for each individual animal/brain or simulation as described above. The subtraction procedure was then performed using custom-written Matlab code, which creates a new null distribution (*k_diff_*) by performing a bootstrapping procedure sampling 90000 times (with replacement) from each distribution and then creates a new null distribution for the differences. *P*-values are computed with the new null distribution by asking how many values in the distribution are equal to or greater than the actual observed difference (note that the smallest *p*-value for an individual comparison is 1/90000 or 1x10^-5^). *P*-values were computed for each pair-wise comparison (i.e. the FF-FR subtraction [n=5 in each group] yields 25 pairwise comparisons) and then combined using a harmonic mean approach. Significance thresholds were set using a stringent Bonferroni correction (α=0.01/620 for FF-FR, RR-FR, and Ghrelin (GG-HH) analyses, and (α=0.05/620) for the semaglutide (SS-HH) and FR-HH analysis.

#### *P*-value Ranking and Visualization

For visual comparison of *p*-values using various methods, we rank-ordered *p*-values obtained for FF-FR TTP-S (as described above) and FF TTP_i_ (without subtraction, as described above) and compared these to *p*-values obtained from t-tests of c-Fos alone (1TP, n=10 fasted mice vs n=4 homecage mice) or raw counts of double-labeled cells (2TP, n=5 FF mice vs n=4 HH mice). The same procedure was performed for refed mice, and comparing fasted with refed mice using 1TP and 2TP methods.

### *t*-Distributed Stochastic Neighbor Embedding (t-SNE) Analysis

The dimensionality and modularity of our whole-brain data was visualized with t-distributed stochastic neighbor embedding (t-SNE) and k-means clustering using custom-written Matlab code. Raw counts of double-labled cells from each brain were combined into matrices for each group (FF, RR, and RF groups). t-SNE was performed using the Barnes-Hut approximation with 3 principal components and was visualized in 2 dimensions. The modularity structure was visualized using k-means clustering limited to 3 clusters. We additionally chose 30 brain areas at random to compare the cluster distribution to that of FF and RR brains.

### Graph Theoretical Network Analysis

Using our TTP-S analyses (see Statistical Methods above), we identified 126 brain areas that reached significance in either FF-FR and RR-FR brains to be used for graph-theoretical network analysis. Pearson-correlation matrices were calculated for the number of double-labeled neurons (corrected for tdT-positive cells) in each brain area across FF (n=5) and RR (n=5) mice. Positive correlation matrices generated and were thresholded at R > 0.6 and negative correlation matrices were thresholded at R< -0.6. The Brain Connectivity Toolbox (Matlab) was used to calculate the following metrics: participation coefficient (PC), within-module degree Z-score (WMDz), degree centrality, eigenvector centrality, and betweenness score. PageRank was additionally calculated independently for positive and negative correlation matrices using Matlab’s built-in Brain Connectivity Toolbox functions. K-means clustering was used to determine the modularity structures of the correlation matrices. Network connectivity was visualized using Gephi software and nodes were positioned using a Force 2 Atlas algorithm.

### Modeling

#### Power Analysis

All modeling simulations were performed using custom-written Matlab code. For the power analysis depicted in Figure 1D, we developed a straightforward model encompassing a brain area with 10000 neurons. Neurons in the network were modeled as Poisson point processes where the spikes occurred with a rate λ (spontaneous activity), and interspike intervals obeyed decaying exponential behavior. Within this network, an engram of 500 neurons, constituting 5 percent of the total, displayed stimulus-driven firing rates represented by λ_s._ We simulated both spontaneous and stimulus induced firing rates over 30 trials and across a range of different spontaneous and stimulus evoked firing rates, simulating data for both one-timepoint (e.g. with c-Fos) and two-timepoint (e.g. tdT double-labeling with c-Fos) screening methods. Sample size calculations were performed using power analysis to determine the number of samples required to detect a significant difference in mean spike counts between spontaneous and stimulus-induced activity for one-timepoint, vs. double-labeling at two-timepoints, vs. TTP_i_. The alpha-value was corrected to account for multiple comparisons assuming 500 total brain areas – i.e. 0.05/500 = 0.0001. For TTP_i_, *p*-values were combined across simulated brain areas using a harmonic mean approach.

#### Brain Area Simulations for Sensitivity

We simulated brain areas in a manner similar to that done in the Power Analysis above, but now with varying total cell counts (from 1000 to 49000) and stimulus-induced activity (from 1 to 30 percent of total neurons). Assuming a large sample size (n=30), the sensitivity for detecting over- and underrepresentation of double-labeled cells compared to chance was calculated for each combination of total cell count and proportion of cells with stimulus-induced activity. A conservative *p*-value of 0.001 was used as the cutoff for sensitivity, and this cutoff was plotted alongside the actual observed values for total cell count and proportion of stimulus-driven cells we observed across n=40 brains (n=5 each for FF, FR, RF, RR, and GG groups; n=4 for HH and SS groups described in this paper; n=7 brains used for a separate study on food cues not presented in the current manuscript). TTP_i_ could successfully detect more double-labeling (or less double-labeling) than expected by chance for the observed brain areas lying to the right of the black curves in Figure 3A,B, so the percentage of brain areas to the right of those curves was used to characterize sensitivity.

#### Brain Area Simulations to Determine Effect of Tonic Activity

We recognized that if a brain area contains a fixed sub-population of neurons with a higher spontaneous firing rate than the remaining neurons in that brain area, TTP_i_ could incorrectly attribute observed double-labeling to stimulus- or event-driven activity at each of two timepoints. We therefore performed simulations to determine the effect of the existence of a sub-population of neurons with a higher spontaneous firing rate on false positives for detecting more double-labeling that would be expected by chance. We varied the percentage of neurons with a higher spontaneous firing rate and the firing rate of those neurons for different combinations of total cell count and proportion of cells with stimulus-evoked activity. Total cell counts (5000 and 20,000) and the proportion of cells with stimulus-evoked activity (0.05 and 0.10) were chosen to be representative of observed data (Fig. 3A,B). The proportion of cells with a higher spontaneous firing rate was varied from 0.0025 to 0.015, with their firing rate varied from 0.2 – 1 in increments of 0.1. For each simulation we then calculated the false positive rate for detecting double-labeling when compared to the null distribution. Simulations were performed over 1000 iterations.

#### Whole Brain Simulation

We conducted a whole-brain simulation to assess neuronal response patterns across multiple brain areas in condition AA or AB; analogous to our fasting (FF) and fast-refed (FR) experimental conditions. The model also incorporated variations in spontaneous neural activity to mimic physiological noise.The simulation was initialized with 500 brain areas with neural parameters randomly assigned to each area by sampling from fixed distributions derived from our empirical data. These parameters included the total cell count in a brain area – 1000, 5000, 10000, or 30000 neurons – the spontaneous firing rate (0.02, 0.04, or 0.1), stimulus-induced firing rate (always 0.6), and the size of the subnetwork (0, 0.05, 0.1, or 0.2 of the total). Tonically active sub-populations were randomly assigned to ∼1/3 of these brain areas (118 in our simulation), and intermingled networks (i.e.-non-overlapping sub-populations activated by stimulus A and B) were also assigned to ∼1/3 of brain areas (116 in our simulation). This resulted in a whole-brain simulation with an assortment of different types of brain areas, where ground truth was predetermined and known. For each brain area, neural activity was simulated under conditions of spontaneous and stimulus-evoked activity, using a Poisson process as described above. Each simulation captured a single timepoint in state A or state B (i.e. fasting vs. refeeding), as well as double-labeling between two independent timepoints in states AA or AB (i.e. – fast-fast vs. fast-refed). Simulations were repeated 5 times to produce a sample size of n=5. We then sought to determine how well each method correctly classifies a brain area responding to state A or state B. Thus, we compared the single timepoint (e.g. c-Fos), two-timepoint (double-labeled cells), TTP_i_, and TTP-S methods for distinguishing between activity elicited by A vs. B. The results of these analyses were then compared to the ground truth. Receiver Operating Characteristic (ROC) curves were generated to evaluate the model’s accuracy in distinguishing between true and false positives concerning neuronal activation patterns across conditions. The area under the curve (AUC) provided a measure of the model’s ability to correctly classify neuronal responses as associated with either state A (e.g., fasting) or state B (e.g., refeeding). Confusion matrices were used to calculate the sensitivity and specificity of the different approaches.

## Author Contributions

The methodology was conceived by AR. Experiments were designed by AR, LR, EJK, CB, AWF, and CDS. Experiments were carried out by AR, LR, EJK, CB, JG, and ER. Software was developed by LAH (BrainJ) and AR (TTP-tools, simulations). Data analysis was done by AR, EJK, CB, and LAH. The paper was written by AR, EJK, AWF, and CDS.

## Funding

Part of this work was supported by the generous support of Boehringer-Ingelheim (CDS, AWF). AR was supported by BBRF-NARSAD Young Investigator Award, NY Obesity and Nutrition Research Center Pilot and Feasibility Grant, and an Early Career Award from NIDDK K08-DK132493. EJK was supported by grants from NIMH (L70-MH134315 & T32-MH015144). ER was supported by an Early Career Award from NIGMS K99-GM153720.

## Acknowledgments

We thank Pia-Kelsey O’Neill and Rahim Hashim for helpful comments and suggestions on the manuscript. We thank members of the Salzman lab and Ferrante lab for helpful comments and suggestions throughout. Imaging for this project was supported by the Zuckerman Institute’s Cellular Imaging platform.

## Notes

### Competing Interest Statement

The authors have declared no competing interest.

