## Supplemental Figures for "Two-timepoint assays of neural responses increase the sensitivity and specificity of single-cell whole-brain activity screens"

**Supplemental Information.**

Figures S1-S7.

Supplemental File S1 – see additional document.

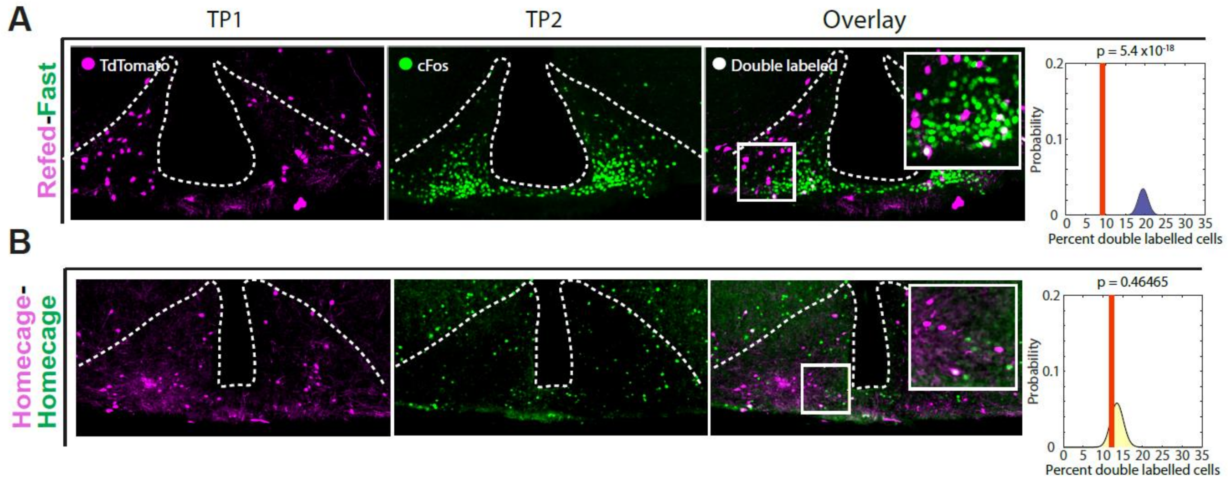


**Figure S1**. **Further examples of TTP analysis from the arcuate nucleus imaged at 10x resolution. A)** A high-resolution image of the arcuate nucleus (ARH) in representative refed-fast (RF) mouse. Timepoint 1 (tdTomato: magenta), timepoint 2 (c-Fos: green), and the overlay with inset showing clear paucity of double-labeled cells. TTP statistical inference ($TTP_i$) plot shown on the right with significant p-value for underrepresentation of double-labeled cells compared to chance. ($p=5.4x10^{-18}$). **B)** Same as A) for an individual HH animal showing minimal double-labeling. Related to Figure 1.

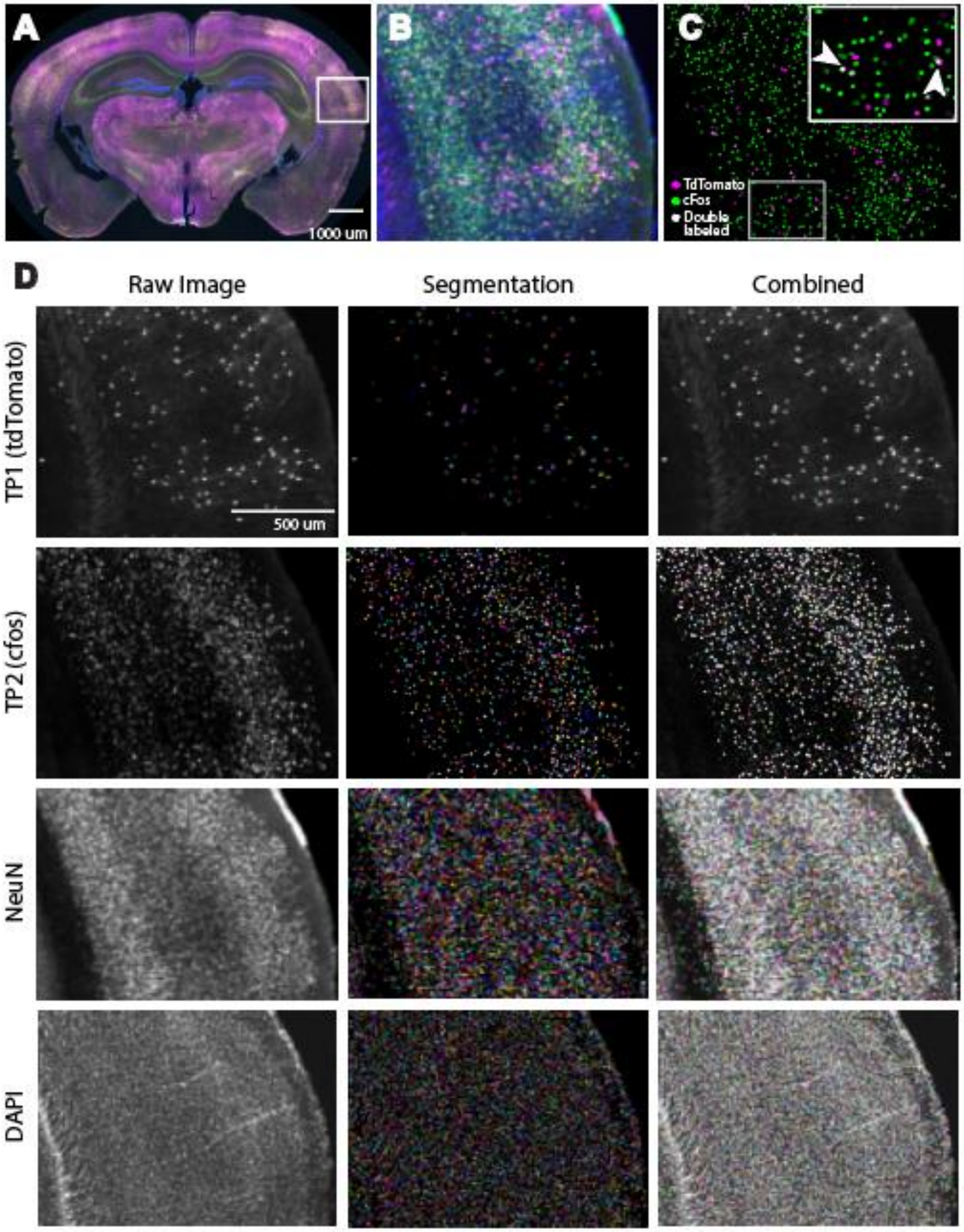


**Figure S2**. **Example segmentation and cell-counting in whole-brain pipeline. A)** An example coronal section taken from whole-brain TTP assay pipeline. **B)** Magnification of boxed area in A**. C)** Segmentations for timepoint 1 (tdTomato; magenta) and timepoint 2 (c-Fos; green), with double-labeled cells (white) illustrated by arrowhead. **D)** Raw image, followed by segmentations, and overlays for the different channels, demonstrating the quality of the segmentations. Related to Figure 2.

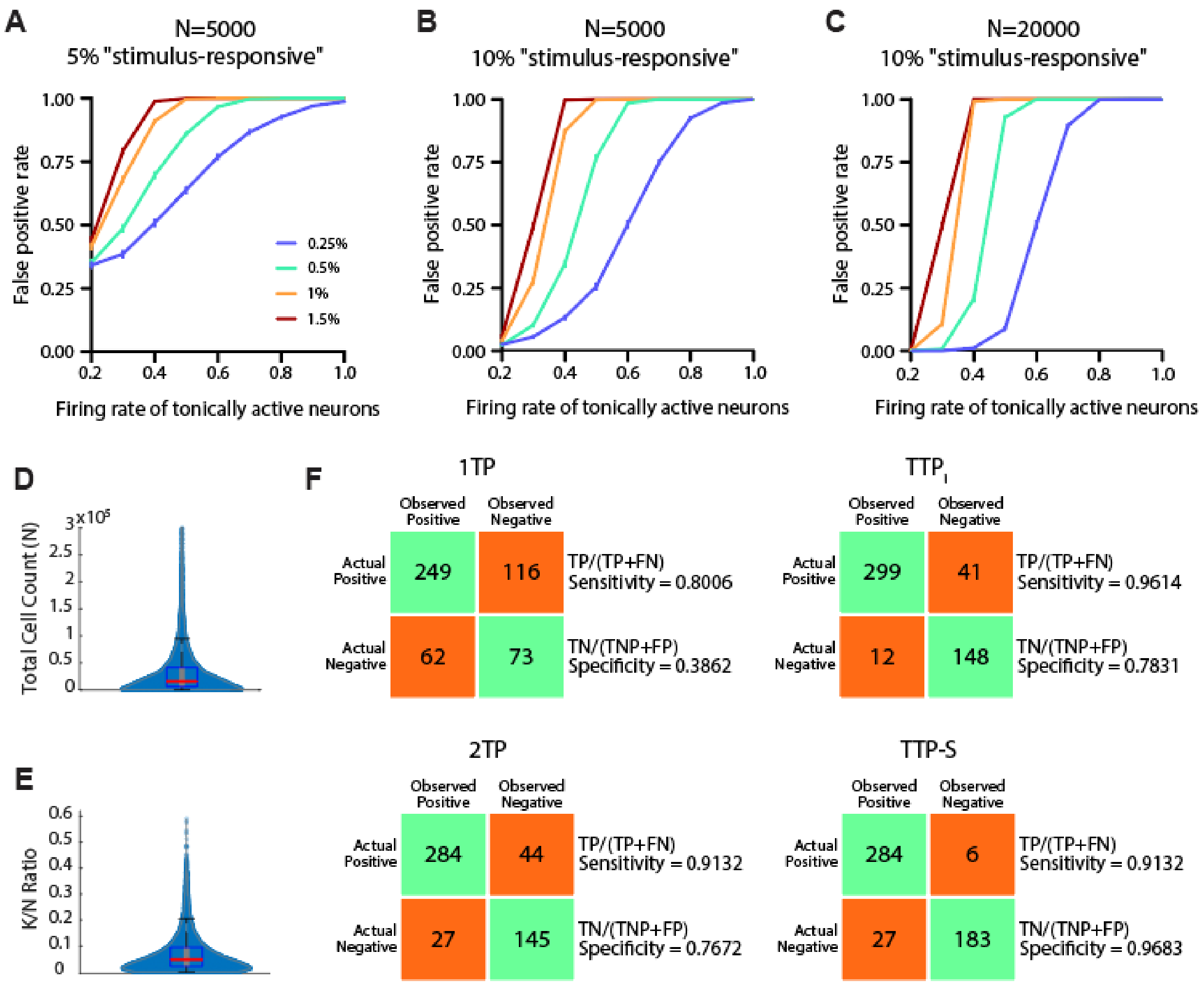


**Figure S3. Further simulation results and cell counts observed in n=40 brains stained in TTP-S pipeline.** Simulated brain region with **A)** total cell count (N) of 5,000, of which 5% are stimulus-responsive; **B)** total cell count (N) of 5,000, of which 10% are stimulus-responsive; and **C)** total cell count (N) of 20,000, of which 10% are stimulus-responsive at both timepoints at an average firing rate of 0.4. The firing rate of 0.25 – 1.5% of separate cells was varied from 0.2 to 1. Increasing the proportion of neurons in the tonically active network as well its firing rate increases the false positive rate. **D)** Violin plot displaying the distribution for total cell count (N) across 22,541 brain areas from n=40 mice. **E)** Violin plot displaying the K/N (c-Fos/total cell count) ratios across 22,541 brain areas from n=40 mice. **F)** Confusion matrices for each method (1TP, analogous to c-Fos staining; 2TP, analogous to raw double-labeled cell counts; $TTP_i$; and TTP-S) in correctly identifying brain regions with stimulus-responsive and intermingled patterns of activity in a simulated whole brain consisting of 500 brain areas. Related to Figure 3.

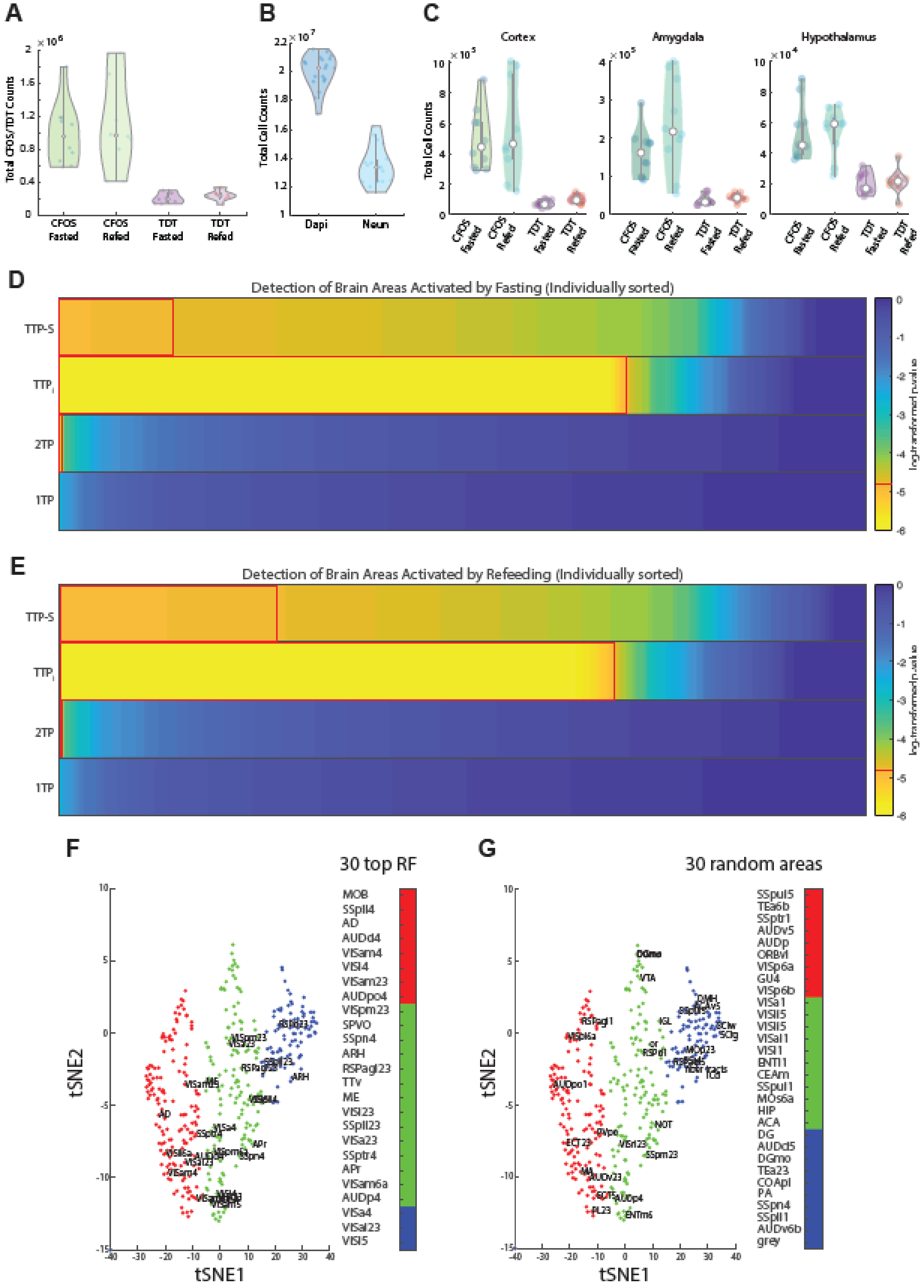

A
Total CFOS/TDT Counts
CFOS Fasted
CFOS Refed
TDT Fasted
TDT Refed
B
Total Cell Counts
Dapi
Neun
C
Cortex
Amygdala
Hypothalamus
Total Cell Counts
CFOS Fasted
CFOS Refed
TDT Fasted
TDT Refed
D
Detection of Brain Areas Activated by Fasting (Individually sorted)
TTP-S
TTP$_i$
2TP
1TP
log-transformed p-value
E
Detection of Brain Areas Activated by Refeeding (Individually sorted)
TTP-S
TTP$_i$
2TP
1TP
log-transformed p-value
F
30 top RF
tSNE2
tSNE1
MOB
SSpll4
AD
AUDd4
VISam4
VISl4
VISam23
AUDpo4
VISpm23
SPVO
SSpn4
ARH
RSPagl23
TTv
ME
VISl23
SSpll23
VISa23
SSptr4
APr
VISam6a
AUDp4
VISa4
VISal23
VISl5
G
30 random areas
tSNE2
tSNE1
SSpul5
TEa6b
SSptr1
AUDv5
AUDp
ORBvl
VISp6a
GU4
VISp6b
VISa1
VISli5
VISli5
VISal1
VISl1
ENTl1
CEAm
SSpul1
MOs6a
HIP
ACA
DG
AUDd5
DGmo
TEa23
COApl
PA
SSpn4
SSpll1
AUDv6b
grey

**Figure S4. Cell counts for TTP-S pipeline and further comparison of methods. A)** c-Fos counts in the fasted (n=10) or refed (n=10) state; and same for tdTomato (tdT). **B)** Total DAPI and NeuN counts across n=20 brains. **C)** c-Fos and TDT counts in different brain regions. **D)** Comparison of ability of 1TP, 2TP, $TTP_i$ and TTP-S to detect brain areas significantly activated during fasting, with 1TP and 2TP methods showing fasted mice (n=10) compared to refed mice (n=10). P-values are shown in a heatmap sorted from lowest-to-highest with significant values ($p<1.6129 \times 10^{-5}$) outlined in red. **E)** Same as D) for detecting brain areas significantly activated by refeeding, with fasted mice as the comparison group for 1TP and 2TP methods. **F)** A tSNE of fast-refed (FR) double-labeled cell counts with 3 clusters color coded. Each dot represents a brain region, and dots that have been labeled are the top 30 most significant areas identified through TTP analysis. Brain area acronyms and color designation are to the right. **G)** Same as F), but 30 random areas are labeled – again illustrating equal distribution across clusters. See Supplemental File S1 for brain region abbreviations.

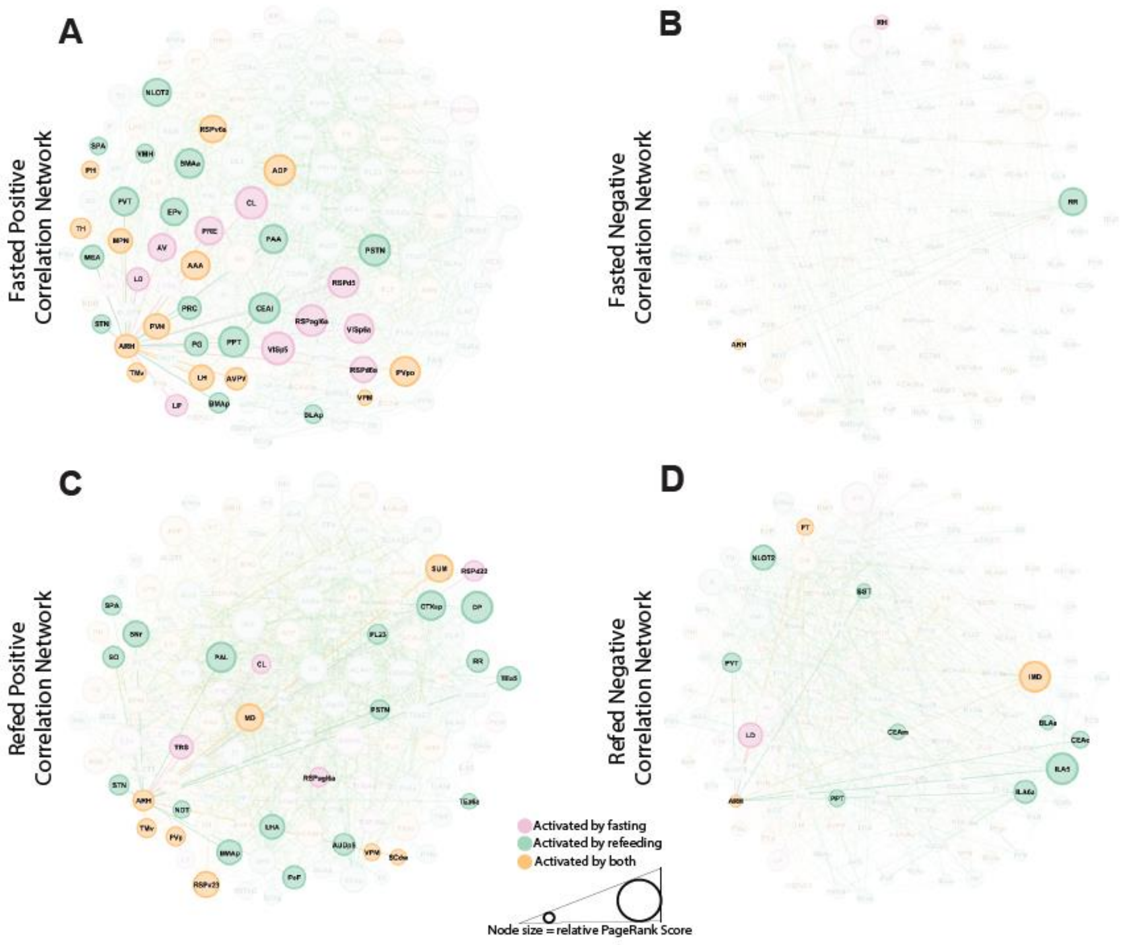


**Figure S5. Graph theoretical analysis to visualize arcuate nucleus functional networks activated during fasting and refeeding.** Graph theoretical approaches were used to visualize the network topology of neurons activated by either the fasted or refed states (126 brain areas). **A**) Positively-correlated network connectivity for the arcuate nucleus (ARH) in the Fasted state (FF mice; n=5). Each line indicates a positive correlation (R>0.6) between brain areas [Fruchterman-Reingold layout]; Node colors indicate whether brain areas were significantly activated by fasting (red/pink), refeeding (green) or both (orange). Only nodes positively correlated with ARH are highlighted. The size of the node indicates the PageRank score, an indication of the connectivity of each node to the network. **B**) Negatively-correlated network connectivity of ARH in the Fasted state (FF mice; n=5). Each line indicates anticorrelated activity (R<-0.6) between brain areas. **C**) Same as A) for ARH but in the refed state (RR mice; n=5). **D**) Same as B) for ARH but in the refed state (RR mice; n=5). Related to Figure 7.

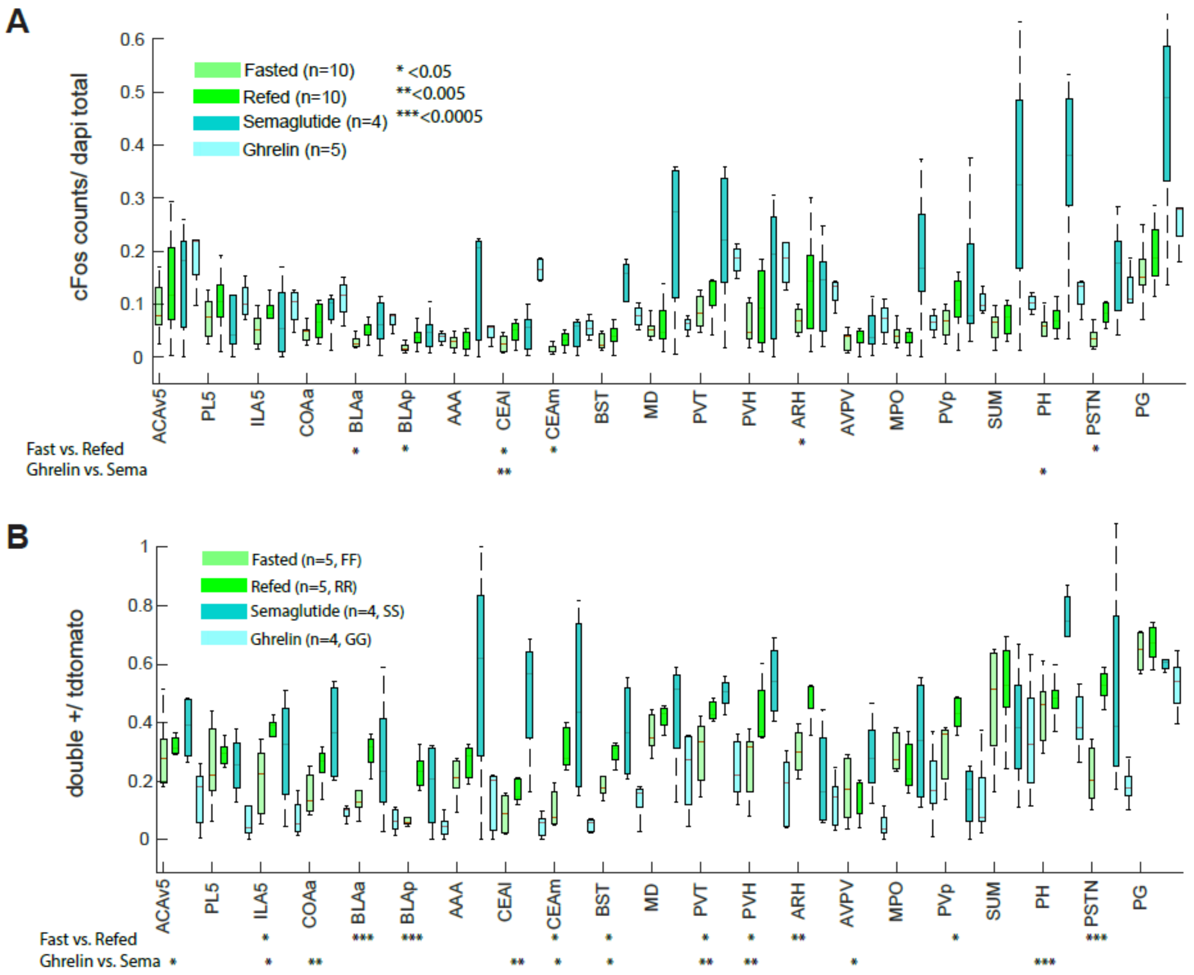


**Figure S6. C-Fos and double-labeled cell counts for selected brain regions identified in semaglutide and ghrelin whole-brain TTP-S analysis. A)** c-Fos counts for the common regions identified between FF-FR and SS-HH mice, as well as RR-FR and GG-HH mice. Plots show c-Fos counts in fasting (n=10) versus refeeding (n=10) versus semaglutide (n=4) versus ghrelin (n=5). P-values below graph indicate raw significance values (not corrected for multiple comparisons). **B)** Same as A), except comparing double-labeled cell counts of fasting (FF; n=5) versus refeeding (RR; n=5), versus semaglutide (SS; n=4) versus ghrelin double-labeled cell (GG; n=5). P-values for t-tests of fasted vs refed mice and semaglutide vs ghrelin mice are displayed under each graph and are not corrected for multiple comparisons. Brain Areas: **ACAv5** – Anterior Cingulate area, ventral part; **PL5** – prelimbic area, layer 5; **ILA5** – infralimbic area, layer 5; **COAa** – cortical amygdalar area, anterior part; **BLAp** – basolateral amygdalar nucleus, posterior part; **BLAa** – basolateral amygdalar nucleus, anterior part; **AAA** – anterior amygdalar area; **CEAl** – Central amygdalar nucleus, lateral part; **CEAm** – Central Amygdalar Nucleus, medial part; **BST** – Bed nuclei of the stria terminalis; **MD** – Mediodorsal nucleus of the thalamus; **PVT** – paraventricular nucleus of the thalamus; **PVH** – paraventricular hypothalamic nucleus; **ARH** – arcuate hypothalamic nucleus; **AVPV** – Anteroventral periventricular nucleus; **MPO** – Medial preoptic area; **PVp** – Periventricular hypothalamic nucleus, posterior part; **SUM** – supramammillary nucleus; **PH** – Posterior hypothalamic nucleus; **PSTN** – parasubthalamic nucleus; **PG** – pontine gray. Related to Figure 8.

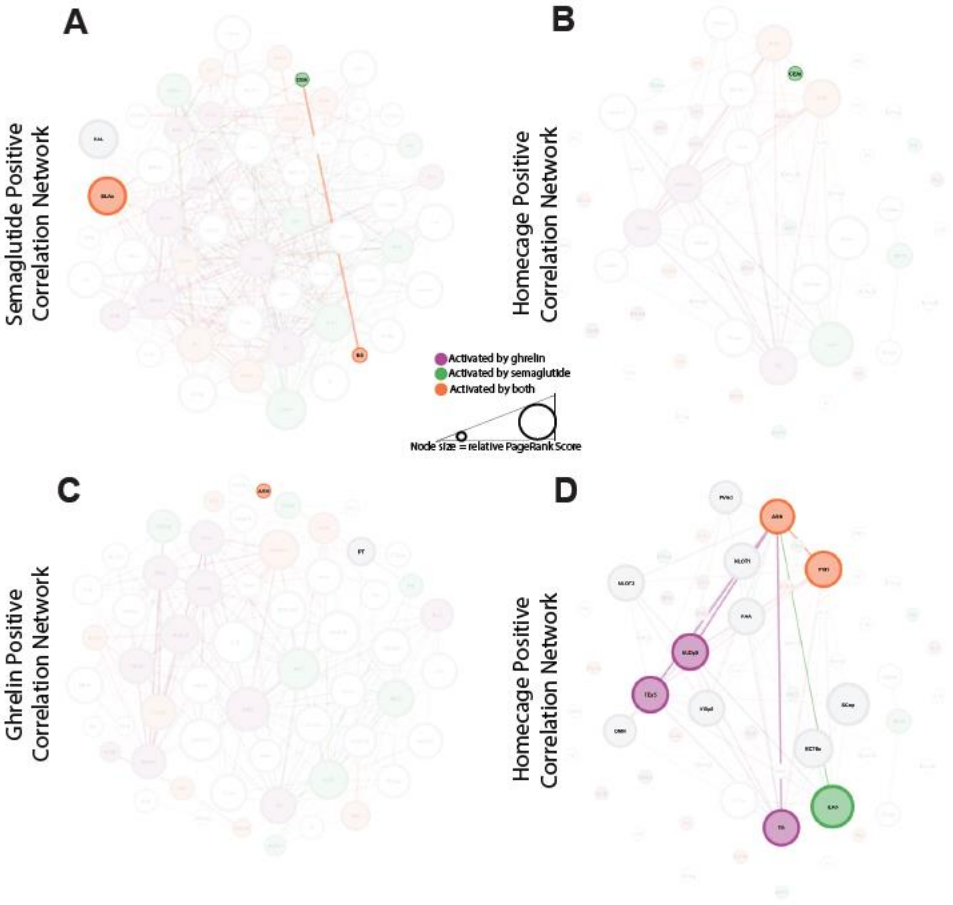


**Figure S7. Positive correlation network plots for semaglutide and ghrelin graph theoretical analyses. A)** Positively-correlated network connectivity for the lateral subdivision of the central amygdala (CEAl) in SS mice (n=4). Each line indicates correlated activity (R<0.6) between brain areas [Fruchterman-Reingold layout]; Node colors indicate whether brain areas were significantly activated by semaglutide (green), ghrelin (purple), or both (orange). Only nodes negatively correlated with CEAl are highlighted. The size of the node indicates the PageRank score, an indication of the connectivity of each node to the network. **B)** Positively-correlated network connectivity for the CEAl in HH control mice (n=4). **C)** Positively-correlated network connectivity for the arcuate nucleus of the hypothalamus (ARH) in GG mice (n=5). **D)** Positively-correlated network connectivity for the arcuate nucleus of the hypothalamus (ARH) in HH control mice (n=4). Related to Figure 8. See Supplemental File S1 for brain region abbreviations.
